# The Geometry of Concept Learning

**DOI:** 10.1101/2021.03.21.436284

**Authors:** Ben Sorscher, Surya Ganguli, Haim Sompolinsky

## Abstract

Understanding the neural basis of the remarkable human cognitive capacity to learn novel concepts from just one or a few sensory experiences constitutes a fundamental problem. We propose a simple, biologically plausible, mathematically tractable, and computationally powerful neural mechanism for few-shot learning of naturalistic concepts. We posit that the concepts that can be learnt from few examples are defined by tightly circumscribed manifolds in the neural firing rate space of higher order sensory areas. We further posit that a single plastic downstream readout neuron learns to discriminate new concepts based on few examples using a simple plasticity rule. We demonstrate the computational power of our proposal by showing it can achieve high few-shot learning accuracy on natural visual concepts using both macaque inferotemporal cortex representations and deep neural network models of these representations, and can even learn novel visual concepts specified only through linguistic descriptors. Moreover, we develop a mathematical theory of few-shot learning that links neurophysiology to behavior by delineating several fundamental and measurable geometric properties of high-dimensional neural representations that can accurately predict the few-shot learning performance of naturalistic concepts across all our numerical simulations. We discuss testable predictions of our theory for psychophysics and neurophysiological experiments.

## 1 Introduction

A hallmark of human intelligence is the remarkable ability to rapidly learn new concepts. Humans can effortlessly learn new visual concepts from only one or a few visual examples^1–4^. We can even acquire visual concepts without any visual examples, by relying on cross-modal transfer from language descriptions to vision. The theoretical basis for how neural circuits can mediate this remarkable capacity for ‘few-shot’ learning of general concepts remains mysterious, despite many years of research in concept learning across philosophy, psychology and neuroscience^5–9^.

Theories of human concept learning are at least as old as Aristotle, who proposed that concepts are represented in the mind by a set of strict definitions^10^. Modern cognitive theories propose that concepts are mentally represented instead by a set of features, learned by exposure to examples of the concept. Two such foundational theories include prototype learning, which posits that features of previously encountered examples are averaged into a set of ‘prototypical’ features^11^, and exemplar learning, which posits that the features of all previously encountered examples are simply stored in memory^5^. However, neither theory suggests how these features might be represented in the brain. In laboratory experiments, these features are either constructed by hand by generating synthetic stimuli which vary along a pre-defined set of latent features^12,13^, or are indirectly inferred from human similarity judgements^14–16^.

In this work, we introduce a theory of concept learning in neural circuits based on the hypothesis that the concepts we can learn from a few examples are defined by tight geometric regions in the space of high-dimensional neural population representations in higher-level sensory brain areas. Indeed, in the case of vision, decades of research have revealed a series of representations of visual stimuli in neural population responses along the ventral visual pathway, including V1, V2, and V4, culminating in a rich object representation in the inferotemporal (IT) cortex^17–19^, allowing a putative downstream neuron to infer the identity of an object based on the pattern of IT activity it elicits^20,21^.

We also hypothesize that sensory representations in IT, acquired through a lifetime of experience, are sufficient to enable rapid learning of novel visual concepts based on just a few examples, without any further plasticity in the representations themselves, by a downstream population of neurons with afferent plastic synapses that integrate a subset of non-plastic IT neural responses.

We test our theory on a diverse array of naturalistic visual concepts using artificial deep neural network (DNN) models of the primate visual hierarchy. DNNs currently constitute the best known models of the primate visual hierarchy, and have been shown to predict neural population responses in V1, V4 and IT^22,23^, the similarity structure of object representations in IT^24^, and human performance at categorizing familiar objects^16,25^. Our approach is corroborated by recent work which demonstrates state-of-the-art performance on few-shot learning benchmarks by training linear classifiers atop fixed DNN representations^26,27^.

Our theory establishes fundamental links between key geometric characteristics of IT-like object representations and the capacity for rapid concept learning. Identifying these geometric features allows us to demonstrate that IT-like representations in trained DNNs are powerful enough to accurately mediate few-shot learning of novel concepts, without any further plasticity, and that this remarkable capacity is due to orchestrated transformations of the geometry of neural activity patterns along the layers of trained DNNs. Furthermore, using neurophysiological recordings in macaques in response to a set of synthetic objects^21^, we demonstrate that neural activity patterns in the primate visual pathway also undergo orchestrated geometric transformations such that few-shot learning performance improves from the retina to V1 to V4 to IT. Intriguingly, despite common patterns of performance along both the macaque ventral visual hierarchy and our current best DNN models of this hierarchy, our theory reveals fundamental differences in neural geometry between primate and DNN hierarchies, thereby providing new targets for improving models.

We further leverage our theory to propose a neurally plausible model of cross-modal transfer in human concept learning, allowing neural representations of linguistic descriptors to inform the visual system about visually novel concepts. Our theory also reveals that long-held intuitions and results about the relative performance of prototype and exemplar learning are completely reversed when moving from low dimensional concepts with many examples, characteristic of most laboratory settings, to high-dimensional naturalistic concepts with very few examples. Finally, we make testable predictions not only about overall performance levels but more importantly about salient *patterns* of errors, as a function of neural population geometry, that can reveal insights into the specific strategies used by humans and non-human primates in concept learning.

## 2 Results

### 2.1 Accurate few-shot learning with non-plastic representations

To evaluate the performance of few-shot learning, we use neural representations of visual concepts derived from a DNN that has been shown to be a good model of object representations in IT cortex^28^ (Fig 1, ResNet50; see Methods 3.5). The DNN is trained to classify 1, 000 naturalistic visual concepts from the ImageNet1k dataset (e.g. ‘dog’, ‘cat’) using millions of images. To study *novel* concept learning, we select a set of 1, 000 new visual concepts, never seen during training, from the ImageNet21k dataset (e.g. ‘coati’, ‘numbat’, Fig. 1**a**; see Methods 3.1). We examine the ability to learn to discriminate between a pair of new concepts, given only a few training images, by learning to classify the activity patterns these training images elicit across IT-like neurons in the feature layer of the DNN (Fig. 1**b**). A particularly simple, biologically plausible classification rule is prototype learning, performed by averaging the activity patterns elicited by the training examples into concept prototypes, 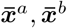 (Fig. 1**b**). A test image is then classified by comparing the activity pattern it elicits to each of the prototypes, and identifying it with the closest prototype. This classification rule can be performed by a single downstream neuron that adjusts its synaptic weights so that its weight vector ***w*** points along the difference between the prototypes, 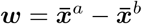 (Fig. 1**b**). Remarkably, prototype learning achieves an average test accuracy of 98.6% across all 1, 000 × 999 pairs of novel concepts given only 5 training images of each concept (Fig. 1**c**), only slightly lower than the accuracy obtained by prototype learning on the same 1,000 familiar concepts that were used to train the DNN (Fig. 1**d**). When only one training example of each concept is provided (1-shot learning), prototype learning achieves a test accuracy of 92.0%. In contrast, the performance of prototype learning applied to representations in the retina-like pixel layer (Fig. 1**b**) or to the feature layer of an untrained, randomly initialized DNN is around 50 − 65% (Fig. 1**c**), indicating that the neural representations learned over the course of training the DNN to classify 1,000 concepts are powerful enough to facilitate highly accurate few-shot learning of novel concepts, without any further plasticity. Consistent with this result, we find that DNNs that perform better on the ImageNet1k classification task also perform better at few-shot learning of novel concepts (Fig. 1**e**). Furthermore, different DNNs are highly consistent in their error patterns (Fig. 1**f**). Interestingly, these error patterns reveal a pronounced asymmetry on many pairs of concepts (SI 6). For instance, models may be much more likely to classify a test example of a ‘coati’ as a ‘numbat’ than a ‘numbat’ as a ‘coati’ (Supp. Fig. 1).

**Figure 1:**
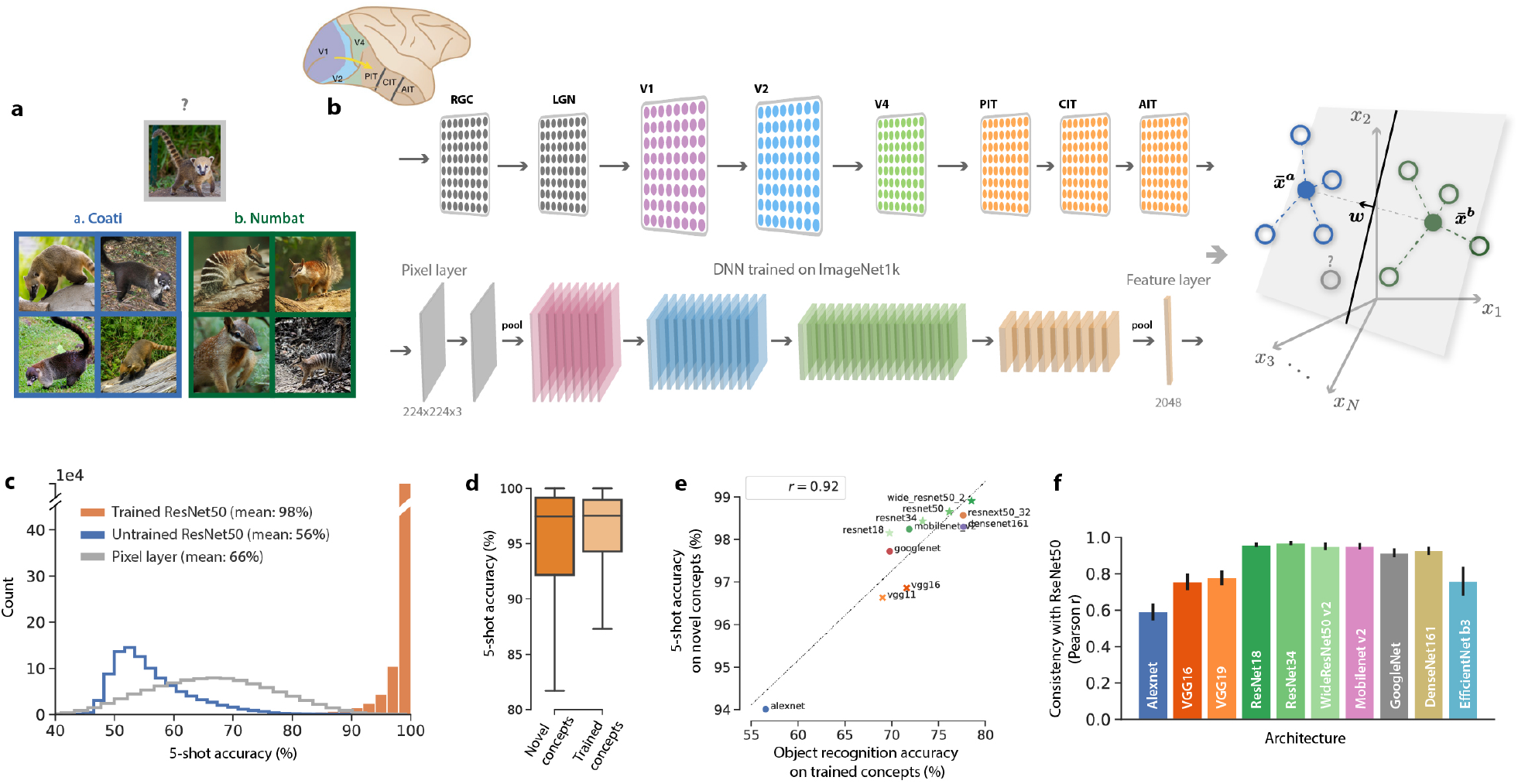
Concept learning framework and model performance. **a**, Example four-shot learning task: does the test image in the gray box contain a ‘coati’ (blue) or a ‘numbat’ (green), given four training examples of each? **b**, Each training example is presented to the ventral visual pathway (top), modeled by a trained DNN (bottom), eliciting a pattern of activity across IT-like neurons in the feature layer. We model concept learning as learning a linear readout ***w*** to classify these activity patterns, which can be thought of as points in a high dimensional activity space (open circles, right). In the case of prototype learning, activity patterns are averaged into prototypes 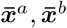 (closed circles), and ***w*** is pointed along the difference between the prototypes 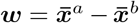, passing through their midpoint. To evaluate generalization accuracy, we present a test image and determine whether its neural representation (gray open circle) is correctly classified. **c**, Generalization accuracy is very high across 1, 000 × 999 pairs of novel visual concepts from the ImageNet21k dataset (orange). In comparison, test accuracy is poor when using a randomly initialized DNN (blue), or when learning a linear classifier in the pixel space of input images (gray). **d**, Test accuracy on novel concepts (dark orange) is only slightly lower than test accuracy on familiar concepts seen during training (light orange). **e**, Performance on the object recognition task used to train DNNs correlates with their ability to generalize to novel concepts given few examples (*r* = 0.92, *p <* 1 × 10^−4^), across a variety of DNN architectures. **f**, Different DNN architectures are consistent in the pattern of errors they make on novel concepts (*p <* 1 × 10^−10^). Error bars are computed by measuring consistency over random subsets of 500 concept pairs.

These results raise several fundamental theoretical questions. Why does a DNN trained on an image classification task also perform so well on few-shot learning? What properties of the derived neural representations empower high few-shot learning performance? Furthermore, why are some concepts easier than others to learn (Fig. 1**c**), and why is the pairwise classification error asymmetric (Supp. Fig. 1)? We answer these questions by introducing an analytical theory that predicts the generalization error of prototype learning, based on the geometry of neural population responses.

### 2.2 A geometric theory of prototype learning

Patterns of activity evoked by examples of any particular concept define a concept manifold (Fig. 2). Although their shapes are likely complex, we find that when concept manifolds are high-dimensional the generalization error of prototype learning can be accurately predicted based only on each manifold’s centroid ***x***_**0**_, and radii *R*_*i*_ along a set of orthonormal basis directions ***u***_*i*_, *i* = 1, …, *N* (see SI 2), capturing the extent of natural variation of examples belonging to the same concept. A useful measure of the overall size of these variations is the mean squared radius 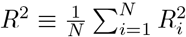.

**Figure 2:**
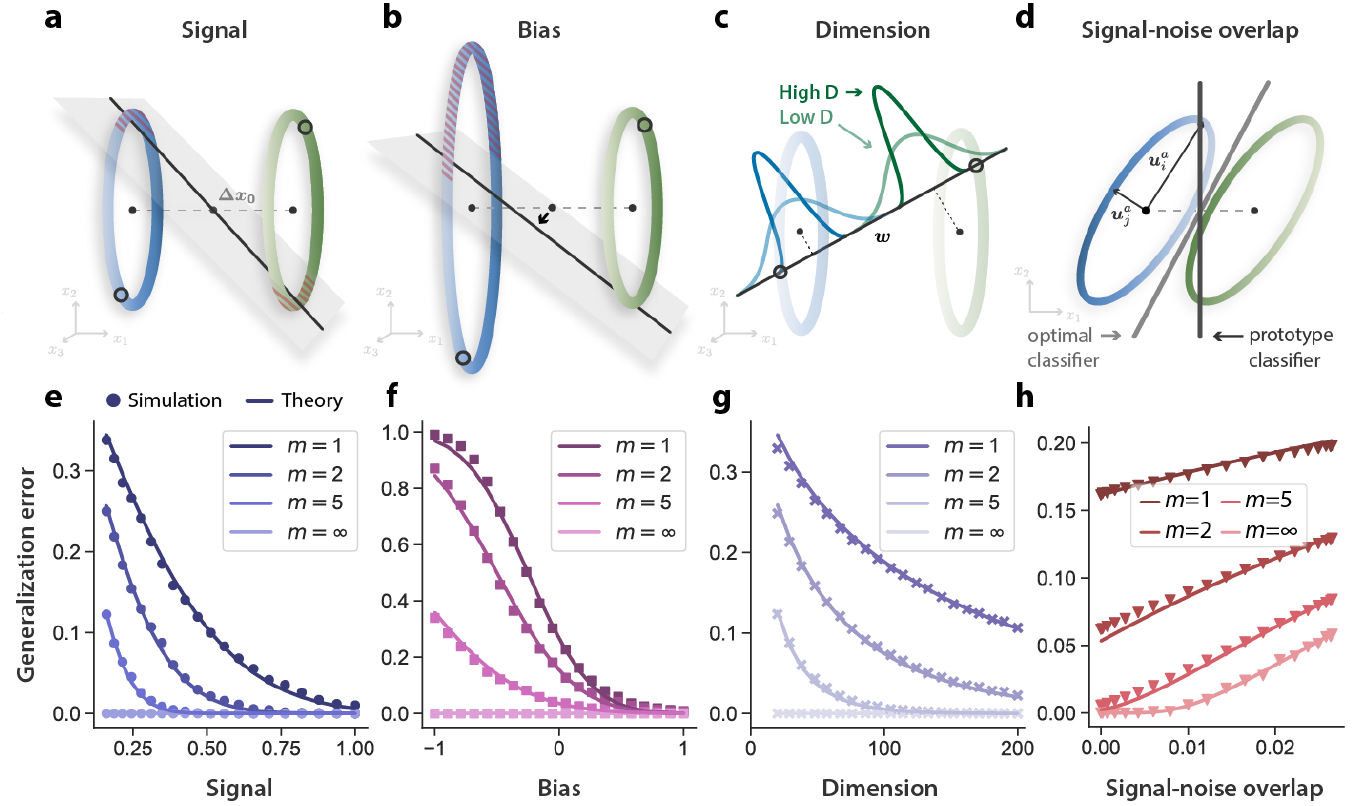
Geometry of few-shot learning. Patterns of activity evoked by examples of each concept define a concept manifold, which we approximate as a high-dimensional ellipsoid. The generalization error (red hashed area) of a prototype classifier ***w*** is governed by four geometric properties of these manifolds (Eq. 1), shown schematically in **a**,**b**,**c**,**d**. Signal, **a**, refers to the pairwise separation between concept manifolds, **‖Δ*x***_**0**_ **‖** ^2^. Manifolds that are better separated are more easily discriminated by a linear classifier (gray hyperplane) given few training examples (open circles). **b**, Bias; as the radius of one manifold grows relative to the other, the decision hyperplane shifts toward the manifold with larger radius. Hence the generalization error on the larger manifold increases, while the error on the smaller manifold decreases. Although the tilt of the decision hyperplane relative to the signal direction averages out over different draws of the training examples, this bias does not. **c**, Dimension; in high dimensions, the projection of each concept manifold onto the linear readout direction ***w*** concentrates around its mean. Hence high-dimensional manifolds are easier to discriminate by a linear readout. **d**, Signal-noise overlap; pairs of manifolds whose noise directions ***u***_*i*_ overlap significantly with the signal direction **Δ*x***_**0**_ have higher generalization error. Even in the limit of infinite training examples, *m*→ ∞, the prototype classifier (dark grey) cannot overcome signal-noise overlaps, as it has access only to the manifold centroids. An optimal linear classifier (light grey), in contrast, can overcome signal-noise overlaps using knowledge of the variability around the manifold centroids. **e**,**f**,**g**,**h**, Behavior of generalization error (with *m* = 1, 2, 5, ∞) in relation to each geometric quantity in **a**,**b**,**c**,**d**. Theoretical predictions are shown as dark lines, and few-shot learning experiments on synthetic ellipsoidal manifolds are shown as points. **f**, When bias is large and negative, generalization error can be greater than chance (0.5). **g**, Generalization error decreases when manifold variability is spread across more dimensions. **h**, In the presence of signal-noise overlaps, generalization error remains nonzero even for arbitrarily large *m*. Details on simulation parameters are given in Methods 3.4.

Our theory of prototype learning to discriminate pairs of new concepts (*a, b*) predicts that the average error of *m*-shot learning on test examples of concept *a* is given by *ε*_*a*_ = *H*(SNR_*a*_), where *H*(·) is the Gaussian tail function 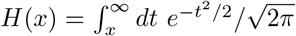(SI 2.3). The quantity SNR_a_ is the signal-to-noise ratio (SNR) for manifold *a*, whose dominant terms are given by,

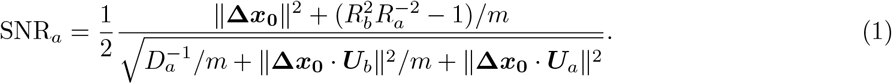

A full expression and derivation are given in SI 2.3. The SNR depends on four interpretable geometric properties, depicted schematically in Fig. 2**a-d**. Their effect on the generalization error is shown in Fig. 2**e-h**. The generalization error for tasks involving discriminating more than two novel concepts, derived in SI 2.4, is governed by the same geometric properties. We now explain each of these properties.

1. **Signal**. 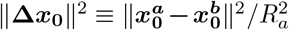 represents the pairwise distance between the manifolds’ centroids, 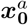 and 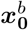, normalized by 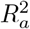 (Fig. 2**a**). As the pairwise distance between manifolds increases, they become easier to separate, leading to higher SNR and lower error (Fig. 2**e**). We denote ‖**Δ*x***_**0**_‖ ^2^ as the signal and **Δ*x***_**0**_ as the signal direction.
2. **Bias**. 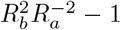 represents the average bias of the linear classifier, (Fig. 2**b**). Importantly, when manifold *a* is larger than manifold *b*, the bias term is negative, predicting a lower SNR for manifold *a*. This asymmetry results from the fact that the classification hyperplane is biased towards the larger manifold (Fig. 2**f**), and causes the asymmetry in generalization error observed above (Sec. 2.1). This bias is unique to few-shot learning. As can be seen in Eq. 1, its effect on SNR diminishes for large numbers of training examples *m*.
3. **Dimension**. In our theory, 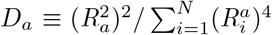, known as the *participation ratio*^29^, quantifies the number of dimensions along which the concept manifold varies significantly, which is often much smaller than the number of neurons *N* (Methods 3.3). Intriguingly, Eq. 1 predicts that higher-dimensional manifolds perform *better* for few-shot learning. This enhanced performance occurs because projections of examples along the readout direction ***w*** concentrate around their mean with variance 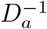 (Fig. 2**c**), hence noise induced by errors in estimating prototypes is suppressed as *D*_*a*_ increases (Fig. 2**g**). Interestingly, this behavior is opposite to that observed in object recognition^30,31^, where a greater number of familiar objects can be perfectly classified if the dimensionality of each manifold is small.
4. **Signal-noise overlap**. ‖**Δ*x***_**0**_·***U***_*a*_‖^2^ and ‖**Δ*x***_**0**_·***U***_*b*_‖ ^2^ quantify the overlap between the signal direction **Δ*x***_**0**_ and the manifold axes of variation 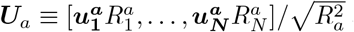 and 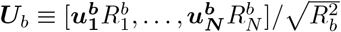 (Fig. 2**d**, and Methods 3.3 for details). Generalization error increases as the overlap between the signal and noise directions increases (Fig. 2**h**). We note that signal-noise overlaps decrease as the dimensionality *D*_*a*_ increases.

#### Effect of number of training examples

As the number of training examples *m* increases, the prototypes more closely match the true manifold centroids; hence the bias and the first two noise terms in Equation 1 decay as 1*/m*. The last noise term however, ‖**Δ*x***_**0**_ ***U***_***a***_ ***‖***^2^, does not vanish even when the centroids are perfectly estimated, as it originates from variability in the *test* examples along the signal direction (Fig. 2**i**, *m* = ∞). Thus, in the limit of a large number of examples *m*, the generalization error does not vanish, even if the manifolds are linearly separable. The SNR instead approaches the finite limit 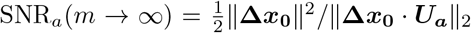, highlighting the failure of prototype learning at large-*m* compared to an optimal linear classifier (Fig. 2**d**). However, in the few-shot (small *m*) regime, prototype learning is close to the optimal linear classifier (see Section 2.8).

To evaluate our theory, we perform prototype learning experiments on synthetic concept manifolds, constructed as high-dimensional ellipsoids. We find good agreement between theory and experiment for the dependence of generalization error on each of the four geometric quantities and on the number of examples (Figure 2**e-h**, and Methods 3.4 for details).

### 2.3 Geometric theory predicts the error of few-shot learning in DNNs

We next test whether our geometric theory accurately predicts few-shot learning performance on naturalistic visual concept manifolds, using the neural representations derived from DNNs studied in Section 2.1. For each pair of concept manifolds, we estimate the four geometric quantities defined above (Methods 3.2), and predict generalization error via Eq. 1, finding excellent agreement between theory and experiment (Fig. 3**a** and Supp. Fig. 3), despite the obviously complex shape of these manifolds.

**Figure 3:**
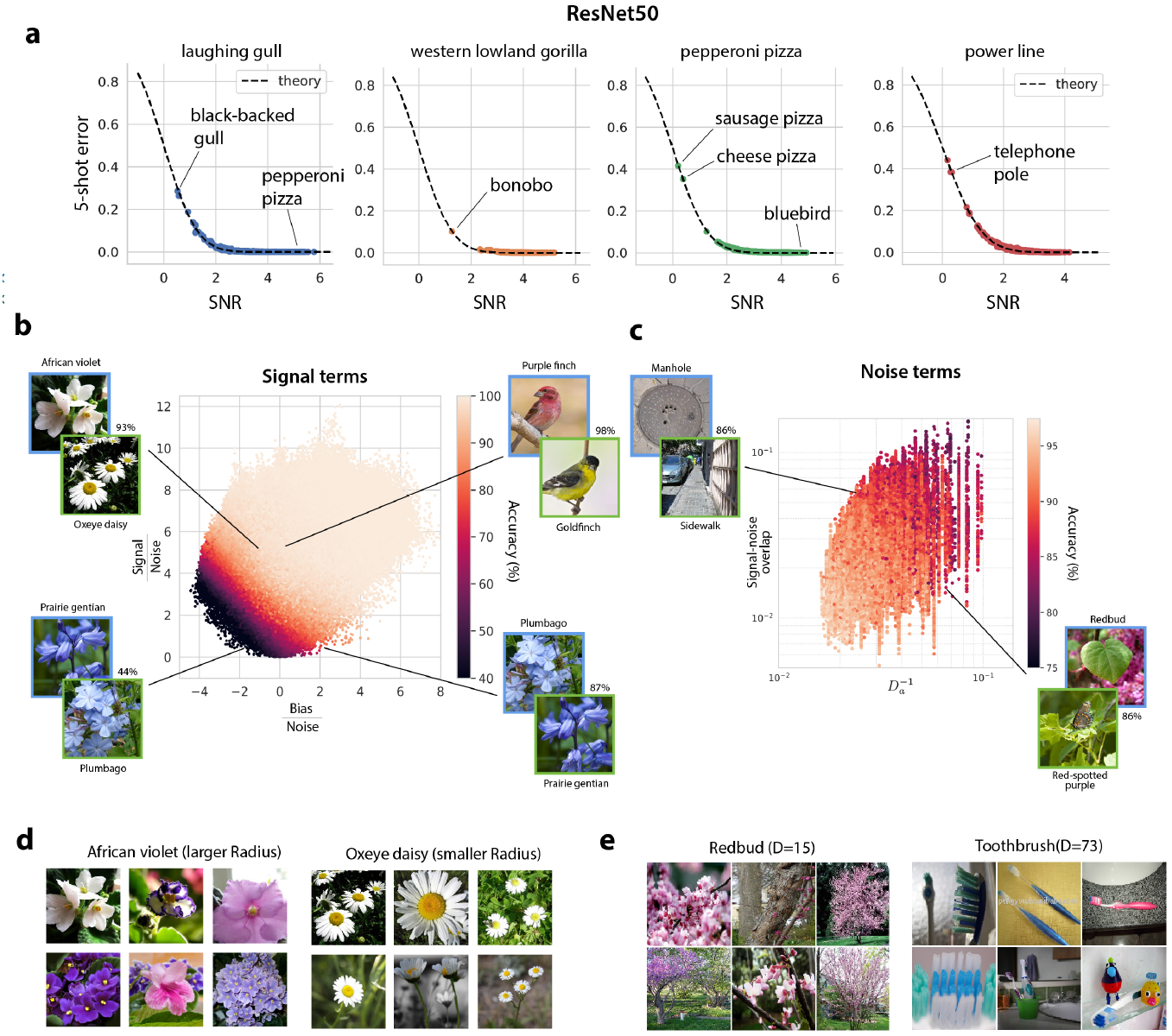
Geometric theory predicts the generalization error of few-shot learning in DNNs. **a**, We compare the predictions of our theory to the few-shot learning experiments performed in Fig. 1. Each panel plots the generalization error of one novel visual concept (e.g. ‘laughing gull’) against all 999 other novel visual concepts (e.g. ‘black-backed gull’, ‘pepperoni pizza’). Each point represents the average generalization error on one such pair of concepts. *x-axis*: SNR (Eq. 1) obtained by estimating neural manifold geometry. *y-axis*: Empirical generalization error measured in few-shot learning experiments. Theoretical prediction (dashed line) shows a good match with experiments. Error bars, computed over many draws of the training and test examples, are smaller than the symbol size. We annotate each panel with specific examples of novel concept pairs, indicating their generalization error. Additional examples are included in Supp. Fig. 3. **b**, Signal terms; we dissect the generalization accuracy on each pair of novel concepts into differential contributions from the signal and bias (Eq. 1). We plot each pair of visual concepts in the resulting signal–bias plane, where both signal and bias are normalized by the noise so that 1-shot learning accuracy (color, dark to light) varies smoothly across the plot. Specific examples of concept pairs are included to highlight the behavior of generalization accuracy with respect to each quantity. For example, the pair “Purple finch” vs “Goldfinch” has a large signal and a bias close to zero, hence a very high 1-shot accuracy (98%). The pair “African violet” vs “Oxeye daisy”, in contrast, has a large signal but a large negative bias; hence its accuracy is lower (93%). Pairs with large negative bias *and* small signal may have very asymmetric generalization accuracy. For instance, “Prairie gentian” vs “Plumbago” has an accuracy of 87%, while “Plumbago” vs “Prairie gentian” has an accuracy of 44%. For each pair of concepts, test examples are drawn from the upper left concept in blue. **c**, Noise terms; we dissect the contributions of dimensionality and signal-noise overlap to generalization error. Because the variability of the signal terms is much larger than that of the noise terms, we include only pairs of concepts whose signal falls within a narrow range, so that we can visually assess whether 1-shot accuracy (color, dark to light) is due to large dimensionality, small signal-noise overlaps, or both. **d**, Visual concepts with larger radius (‘African violet’) exhibit more variation in their visual features than do concepts with smaller radius (‘Oxeye daisy’). **e**, We include an example of a low-dimensional concept manifold (‘Redbud’, *D* = 7), and a high-dimensional concept manifold (‘Toothbrush’, *D* = 73). See Supp. Fig. 6 for further examples.

Our theory further allows us to dissect the specific contribution of each of the four geometric properties of the concept manifolds to the SNR, elucidating whether errors arise from small signal, negative bias, low dimension, or large signal-noise overlap. We dissect the contributions of signal and bias in Fig. 3**b**, and the contributions of dimension and signal-noise overlap in Fig. 3**c**, along with specific illustrative examples.

Interestingly, we observe that manifolds with large radii exhibit much more variation in color, shape, and texture than manifolds with small radii (Fig. 3**d**, and Supp. Fig. 6). Moreover, individual visual concept manifolds in the trained DNN occupy only a small number, ∼ 35, of the 2,048 available dimensions in the feature layer (examples shown in Fig. 3**e**, and Supp. Fig. 6). Finally, we find that the geometry of concept manifolds in the DNN encodes a rich semantic structure, including a hierarchical organization, which reflects the hierarchical organization of visual concepts in the ImageNet dataset (SI 6). As a consequence, pairs of nearby concepts on the semantic tree are more difficult to learn than pairs of distant concepts are (Supp. Fig. 1).

### 2.4 Concept learning along the visual hierarchy

How the ventral visual hierarchy converts low-level retinal representations into higher-level IT representations useful for downstream tasks constitutes a fundamental question in neuroscience. We first examine this transformation in models of the ventral visual hierarchy and later compare to the macaque hierarchy. Our theory enables us to not only investigate the performance of few-shot concept learning along successive layers of the visual hierarchy, but also obtain a finer-resolution decomposition of this performance into the geometric properties of the concept manifolds in each layer. The generalization error of few-shot prototype learning decreases consistently with layer depth across three different trained network architectures (Fig. 4**a**). In contrast, for an untrained network the error increases with depth. Consistent with the decrease in error, SNR increases with depth in all three networks (Fig. 4**b**). Interestingly, the constituent geometric quantities exhibit more subtle, non-monotonic behavior across layers (Fig. 4**c**,**d**,**e**,**f**). Dimension *D* increases by nearly a factor of ten in the early layers and drops back down in the final layers, similar to the behavior observed in a recent work using different dimensionality metrics^32^. Notably, both noise terms, *D*^−1^ and signal-noise overlap, are suppressed by over an order of magnitude in the late layers of the trained DNNs relative to the untrained DNN.

**Figure 4:**
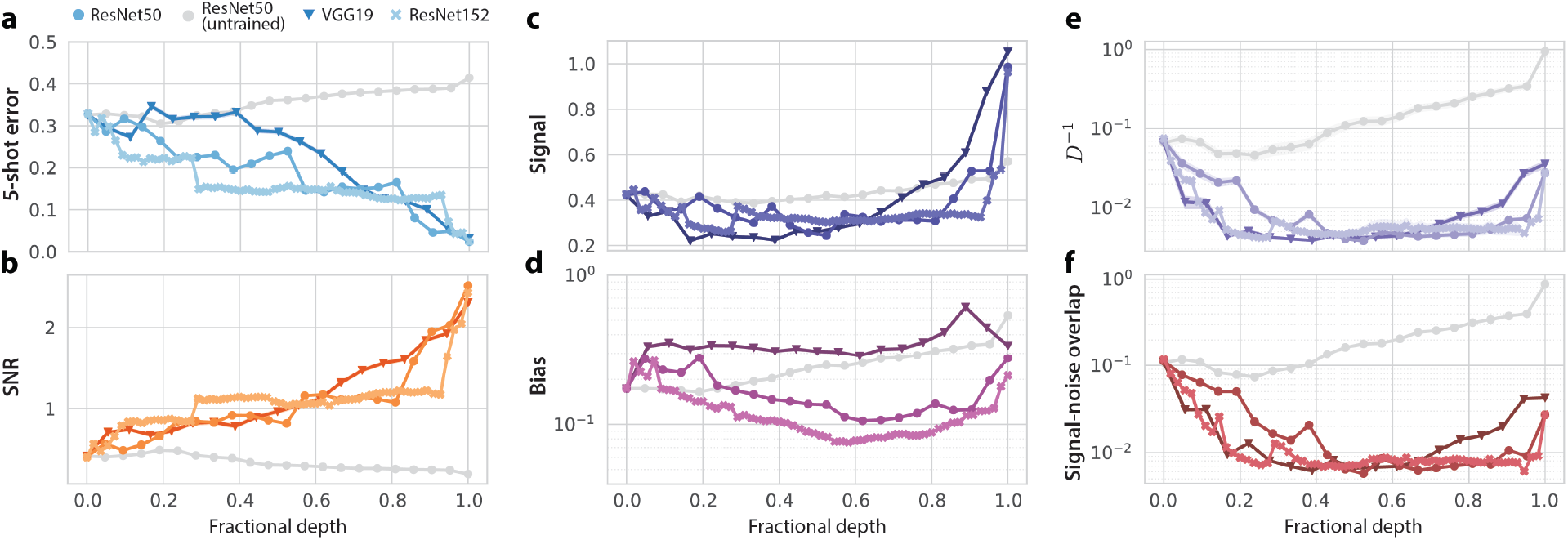
Few-shot learning improves along the layers of a trained DNN, due to orchestrated transformations of concept manifold geometry. Each panel shows the layerwise behavior of one quantity in three different trained DNNs, as well as an untrained ResNet50 (grey). Lines and markers indicate mean value over 100 × 99 pairs of objects; surrounding shaded regions indicate 95% confidence intervals (often smaller than the symbol size). **a**, Few-shot generalization error decreases, and **b**, SNR increases roughly monotonically along the layers of each trained DNN. Signal, **c**, increases dramatically in the later layers of the DNN. Bias (**d**) does not change significantly from pixel layer to feature layer. Noise terms: 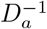 (**e**) and signal-noise overlap (**f**) decrease from pixel layer to feature layer, first decreasing sharply in the early layers of the network, and then increasing slightly in the later layers of the network. Both 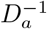 and signal-noise overlap in the trained DNNs are more than on order of magnitude smaller than in their untrained counterpart, indicating that a prominent effect of training is to suppress these noise terms, even though neither noise term is explicitly penalized in the classification objective used to train the DNNs.

### 2.5 Concept learning using primate neural representations

We next investigate the geometry of concept manifolds in the primate visual hierarchy, obtained via recordings of macaque V4 and IT in response to 64 synthetic visual concepts, designed to mimic naturalistic stimuli^21^. We use our geometric theory to predict the generalization error of few-shot prototype learning experiments on these concept manifolds. The results in Fig. 5**a** show an excellent agreement between theory and experiment, and an average 5-shot accuracy of 84% across all 64 × 63 pairs of visual concepts.

**Figure 5:**
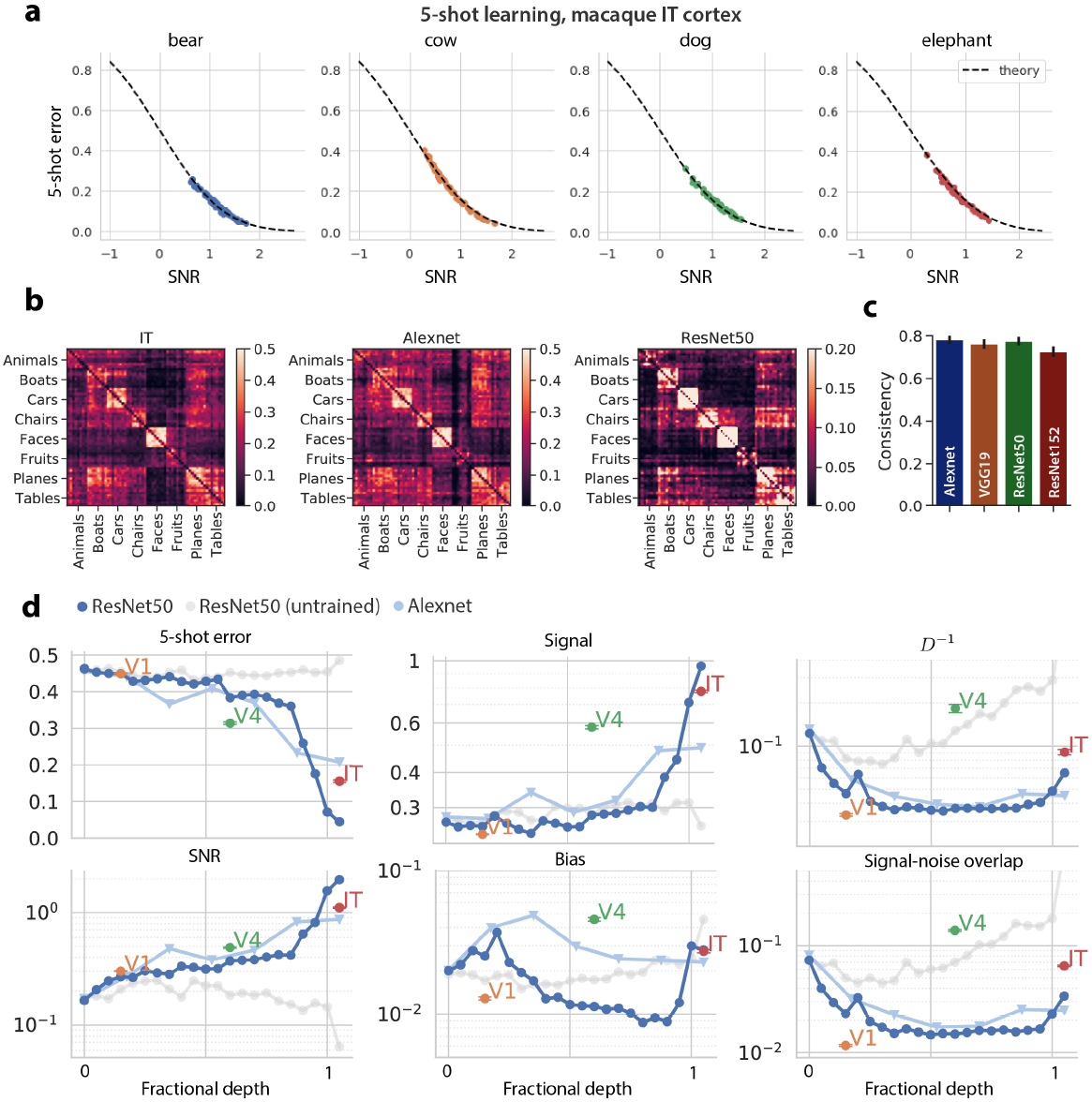
Concept learning and manifold geometry in macaque IT. **a**, 5-shot prototype learning experiments performed on neural population responses of 168 recorded neurons in IT^21^. Each panel shows the generalization error of one visual concept (e.g. ‘bear’) against all 63 other visual concepts (e.g. ‘cow’, ‘dog’). Each point represents the average generalization error on one such pair of concepts. *x-axis:* SNR (eq. 1) obtained by estimating manifold geometry. *y-axis:* Empirical generalization error measured in few-shot learning experiments. The result shows a good fit to the theory (dashed line). Error bars, computed over many draws of the training and test examples, are smaller than the symbol size. **b**, Error pattern of 5-shot learning in IT reveals a clear block diagonal structure, similar to the error patterns in AlexNet and ResNet50. **c**, Error patterns in DNNs are consistent with those in IT (Pearson *r, p <* 1 × 10^−10^). Error bars are computed by measuring consistency over random subsets of 500 concept pairs. **d**, Few-shot learning improves along the ventral visual hierarchy from pixels to V1 to V4 to IT, due to orchestrated transformations of concept manifold geometry. We compute the geometry of concept manifolds in IT, V4, and a simulated population of V1 neurons (see Methods 2.5 for details). Each panel shows the evolution of one geometric quantity along the visual hierarchy. The layerwise behavior of a trained ResNet50 (blue), Alexnet (light blue), and an untrained ResNet50 (grey) is included for comparison. We align V1, V4, and IT to the ResNet layer which best predicts population activity in each cortical area via linear regression, using the BrainScore metric^28^ (see Methods 2.5 for details). Overall performance and SNR exhibit a close match between the primate visual pathway and trained DNNs, while individual geometric quantities display a marked difference. Signal is significantly larger in V4 than in the corresponding DNN layer. Furthermore, both noise terms (*D*^−1^ and signal-noise overlaps) are significantly higher in V4 than in the corresponding DNN layer. These effects trade off so that the overall SNR (and hence few-shot performance) yields a close match between the primate visual hierarchy and the trained DNN. Lines and markers indicate mean value over 64 × 63 pairs of concepts; surrounding error bars and shaded regions indicate 95% confidence intervals.

This performance is slightly better than that achieved by the AlexNet DNN (80%) on the same set of visual concepts, and worse than ResNet50 (93%), consistent with previous experiments on object recognition performance in primates^21^. We predict that performance based on IT neurons will improve when evaluated on more naturalistic visual stimuli than the grayscale, synthetic images used here (as suggested by the increased performance of DNNs, e.g. from 80% to 94% for AlexNet when using novel stimuli from ImageNet).

Beyond overall performance, we find that the error *patterns* in IT and DNNs are strikingly similar (Fig. 5**b**), all exhibiting a prominent block diagonal structure, reflecting the semantic structure of the visual concepts^21^. All networks tested were highly consistent with IT in few-shot learning performance across concept pairs (Fig. 5**c**, *p <* 1 × 10^−10^).

To examine the evolution of few-shot learning capability across the cortical hierarchy, we compute the SNR in IT, V4, and a simulated population of V1 neurons (Methods 3.6). We find that the SNR, and hence few-shot learning performance, increases along the visual hierarchy from just above chance in the pixel layer and V1 to 69% in V4, and 84% in IT, approximately matching the corresponding layers in trained DNNs (Fig. 5**d**). Interestingly, when we decompose the SNR into its finer-grained geometric components, we find that the signal, dimension, and signal-noise overlaps in the primate visual cortex depart substantially from their layerwise behavior in the trained DNN – particularly in V4, where manifold dimension is nearly 10 times lower than in the corresponding DNN layer, while signal is several times larger. The increased noise and increase signal balance out, so that overall SNR and performance is still similar.

### 2.6 How many neurons are required for concept learning?

Until now we have assumed that a downstream cortical neuron has access to the entire neural representation. Can a more realistic neuron which only receives inputs from a small fraction of IT-like neurons still perform accurate few-shot learning? Similarly, can a neuroscientist who only records a few hundred neurons reliably estimate the geometry of concept manifolds (as we have attempted above)?

Here we answer these questions by drawing on the theory of random projections^33–35^ to estimate the effect of subsampling a small number of *M* neurons (see SI 4 for details). We find that subsampling causes distortions in the manifold geometry that decrease both the SNR and the estimated dimenstionality, as a function of the number of recorded neurons *M*,

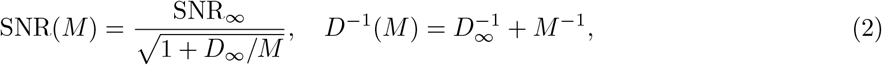

where SNR_*∞*_ and *D*_*∞*_ are the asymptotic SNR and dimensionality given access to arbitrarily many neurons (see SI 4). However, these distortions are negligible when *M* is large compared to the asymptotic dimensionality *D*_*∞*_. Indeed, in both macaque IT and a trained DNN model (Fig. 6), a downstream neuron receiving inputs from only about 200 neurons performs essentially similarly to a downstream neuron receiving inputs from all available neurons (Fig. 6**a**,**b**), and with recordings of about 200 IT neurons the estimated dimensionality approaches its asymptotic value (Fig. 6**c**,**d**; *D*_*∞*_ *≈* 35 in the trained DNN and *D*_*∞*_ *≈* 12 in macaque IT).

**Figure 6:**
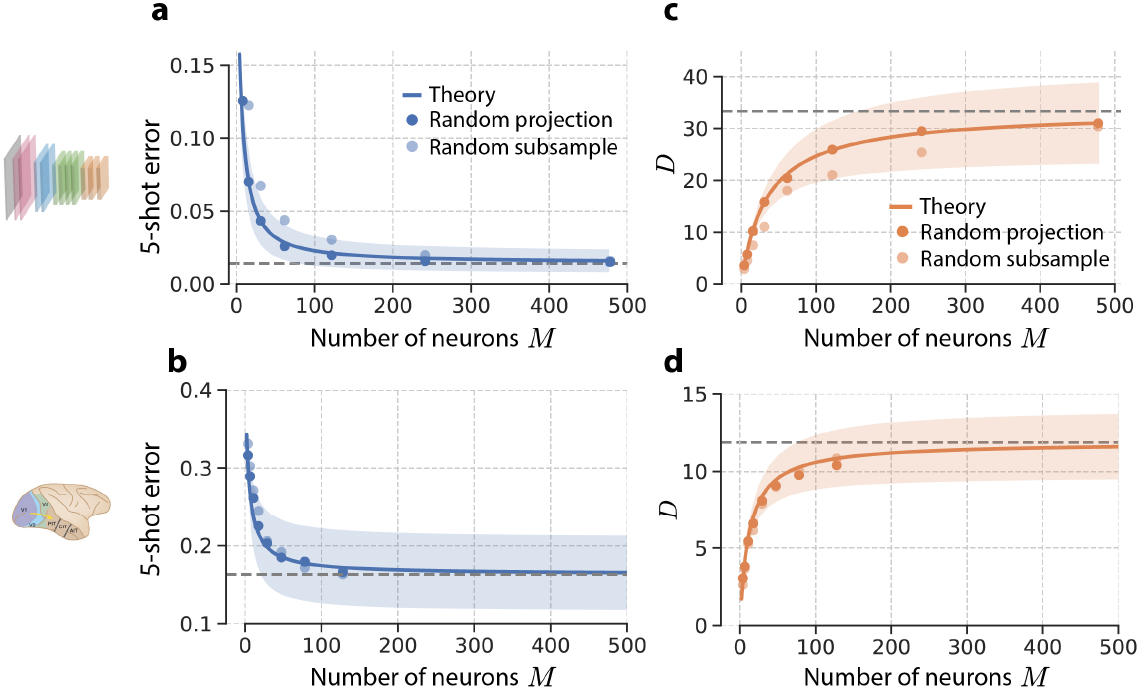
Effect of number of sampled neurons on concept learning and manifold geometry. 5-shot learning experiments in (**a**) ResNet50 on 1, 000 × 999 pairs of concepts from the ImageNet21k dataset and in (**b**) macaque IT on 64 × 63 pairs of novel visual concepts^21^, given access to only *M* neurons (light blue points), and given access to only *M* random linear combinations of the *N* available neurons (dark blue points). The blue curve represents the prediction from the theory of random projections, Eq. 2, the dashed line is its predicted asymptotic value, SNR_*∞*_, and the shaded region represents the standard deviation over all pairs of concepts. In each case, the 1-shot learning error remains close to its asymptotic value provided the number of recorded neurons *M* is large compared to the asymptotic manifold dimension *D*_*∞*_. **c**,**d**, The estimated manifold dimension *D*(*M*) as a function of *M* randomly sampled neurons (light orange points), and *M* random linear combinations (dark orange points) of the *N* neurons in (**c**) the ResNet50 feature layer and in (**d**) macaque IT. The orange curve represents the prediction from the theory of random projections, the dashed line is its predicted asymptotic value, *D*_*∞*_, and the shaded region represents the standard deviation over all pairs of concepts.

### 2.7 Visual concept learning without visual examples

Humans also possess the remarkable ability to learn new visual concepts using only linguistic descriptions, a phenomenon known as *z* ero-shot learning (Fig. 7**a)**. The rich semantic structure encoded in the geometry of visual concept manifolds (Supp. Fig. 1) suggests that a simple neural mechanism might underlie this capacity, namely learning visual concept prototypes from *non-visual* language representations. To test this hypothesis, we obtain language representations for the names of the 1,000 familiar visual concepts used to train our DNN, from a standard word vector embedding model^36^ trained to produce a neural representation of all words in English based on co-occurrence statistics in a large corpus (see Methods 3.7).

**Figure 7:**
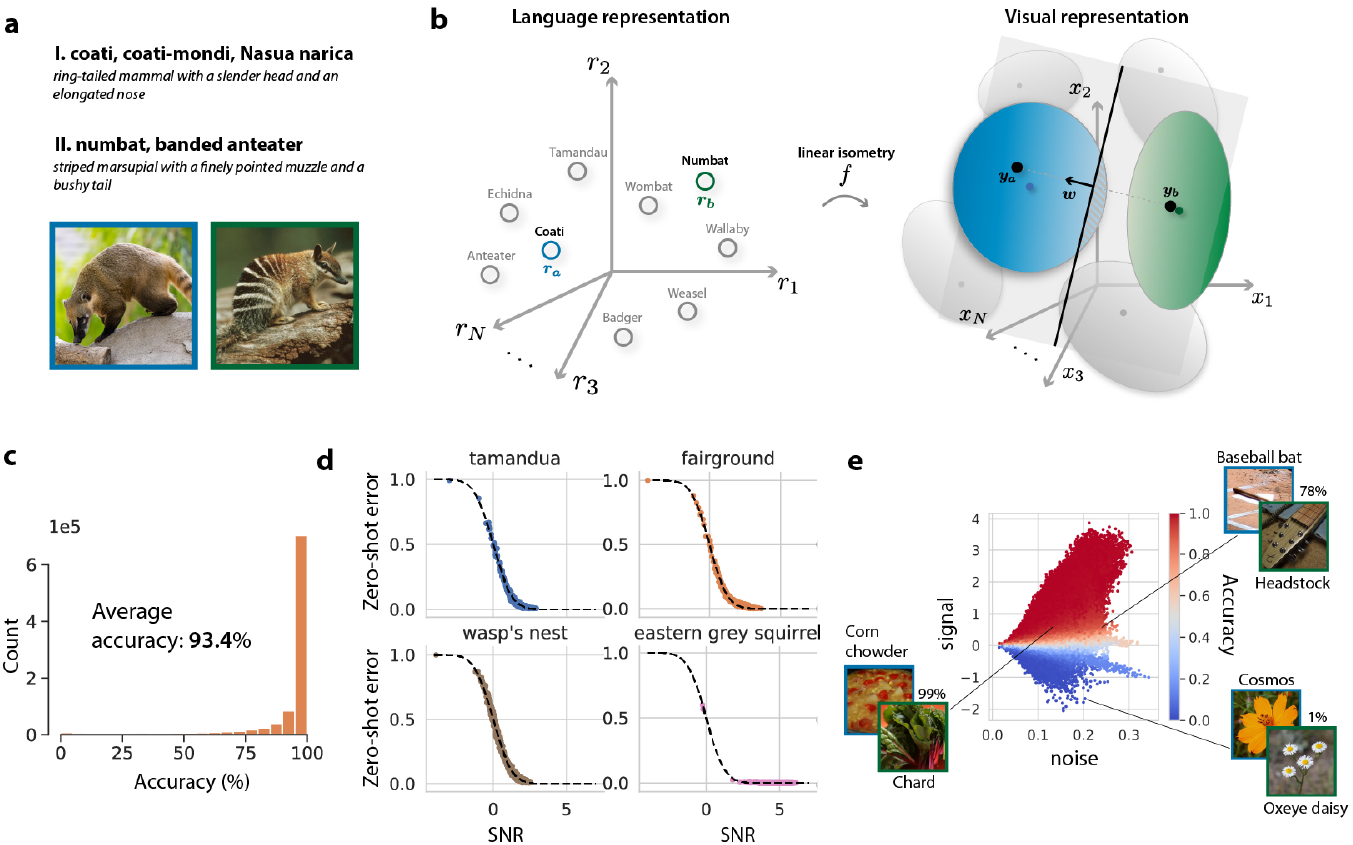
Visual concept learning without visual examples. **a**, Example of the task of learning novel visual concepts given only language descriptions (zero-shot learning): can you identify which image contains the ‘coati’, and which the ‘numbat’? **b**, To simulate this task, we collect language representations from a word vector embedding model (left, gray circles) for the names of the 1, 000 visual concepts used to train the DNN (e.g. ‘anteater’, ‘badger’), along with corresponding visual concept manifolds from the trained DNN (right, gray ellipsoids). We learn a linear isometry *f* to map each language representation as closely as possible to its corresponding visual manifold centroid (Methods 3.7). To learn novel visual concepts (e.g. ‘coati’, and ‘numbat’) without visual examples, we obtain their language representations ***r***_*a*_ from the language model and map them into the visual representation space via the linear isometry, ***y***_*a*_ = *f* (***r***_*a*_). Treating the resulting representations ***y***_*a*_ as visual prototypes, we classify pairs of novel concepts by learning a linear classifier ***w*** (grey hyperplane) which points along the difference between the prototypes, ***w*** = ***y***_*a*_ − ***y***_*b*_, passing through their midpoint. Generalization error (red hashed area) is evaluated by passing test images into the DNN and assessing whether their visual representations are correctly classified. **c**, Generalization accuracy is high across all 1, 000 × 999 pairs of novel visual concepts. **d**, Each panel shows the generalization error of one visual concept against the 999 others. Each point represents the average generalization error on one such pair of concepts. *x-axis*: SNR (Eq. 3) obtained by estimating neural manifold geometry. *y-axis*: Empirical generalization error measured in zero-shot learning experiments. Theoretical prediction (dashed line) shows a good match with experiments. Error bars, computed over many draws of the training and test examples, are smaller than the symbol size. **e**, We decompose the SNR into contributions from the signal and the noise (Eq. 3), and plot each pair of concepts in the resulting signal-noise plane. Zero-shot learning accuracy (blue: low, red: high) varies smoothly across the plot. We annotate the plot with a few representative examples. Some visual concepts have such poor language-derived prototypes that the zero-shot learning accuracy is worse than chance (e.g. on the pair of flowers: ‘cosmos’ and ‘oxeye daisy’, accuracy is only 1%). Color varies predominantly along the signal direction, indicating that much of the variation in zero-shot learning performance is due to variation in the quality of language-derived prototypes, in terms of how closely they match their corresponding visual concept manifold centroids (see Supp. Fig. 4).

We then learn a mapping between the language and vision domains (Fig. 7**b**), an approach studied in previous works^37,38^. However, unlike previous works which jointly optimize language and vision representations to match as closely as possible, our language and vision representations are optimized on entirely independent objectives (word co-occurrence and object recognition). Yet remarkably, we find that the language and vision representations can be aligned by a simple linear isometry (rotation, translation, and overall scaling). Furthermore, this alignment generalizes to *novel* concepts, allowing us to construct prototypes for novel visual concepts by simply passing their names into the word vector embedding model and applying the linear isometry. We use these language-derived prototypes to classify visual stimuli from pairs of novel visual concepts, achieving a 93.4% test accuracy (Fig. 7**c**). Intriguingly, this performance is slightly *better* than the performance of 1-shot learning (92.0%), indicating that the name of a concept can provide at least as much information as a single visual example, for the purpose of classifying novel visual concepts (see Supp. Fig. 4).

Our geometric theory for few-shot learning extends naturally to zero-shot learning, allowing us to derive an analogous zero-shot learning SNR (SI 3),

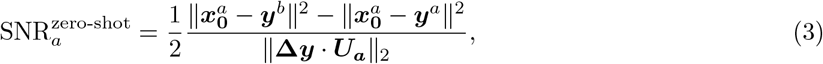

where ***y***^*a*^, ***y***^*b*^ are the language-derived prototypes, and 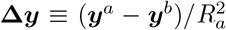 is the associated signal. Fig. 7**d** indicates a close match between this theory and zero-shot experiments.

A surprising prediction of Eq. 3 is that if the distance between the language-derived prototype ***y***^*a*^ and the true visual prototype 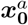 is very large, the SNR may become *negative*, yielding a generalization error worse than chance (examples in Fig. 7**e**) analogous to the bias in Eq. 1.

### 2.8 Comparing cognitive learning models on naturalistic tasks

Our paradigm of using high-dimensional neural representations of naturalistic concepts as features for few-shot learning presents a unique opportunity to compare different concept learning models, such as prototype and exemplar learning, in an ethologically relevant setting, and explore their joint dependence on concept dimensionality *D* and the number of examples *m*. We formalize exemplar learning by a nearest-neighbor decision rule (NN, see Supp. Fig. 8) which stores in memory the neural representation of all 2*m* training examples and classifies a test image according to the class membership of the nearest training example. We find that exemplar learning outperforms prototype learning when evaluated on low-dimensional concept manifolds with many training examples, consistent with past psychophysics experiments on low-dimensional artificial concepts^12,39–41^ (Fig. 8**a**, Methods 3.8). However, prototype learning outperforms exemplar learning for high-dimensional naturalistic concepts and few training examples. In fact, to outperform prototype learning, exemplar learning requires a number of training examples exponential in the number of dimensions, *m* ∼ exp(*D/D*_0_) for some constant *D*_0_ (SI 5). Hence for high-dimensional manifolds like those in trained DNNs and macaque IT, prototype learning outperforms exemplar learning given few examples.

A well-known criticism of prototype learning is that averaging the training examples into a single prototype may cause the prototype learner to misclassify some of the training examples themselves^5^. However, this phenomenon relies on training examples overlapping significantly along a given direction, and almost never happens in high dimensions (Fig. 8**b**) where training examples are approximately orthogonal (Fig. 8**c**).

**Figure 8:**
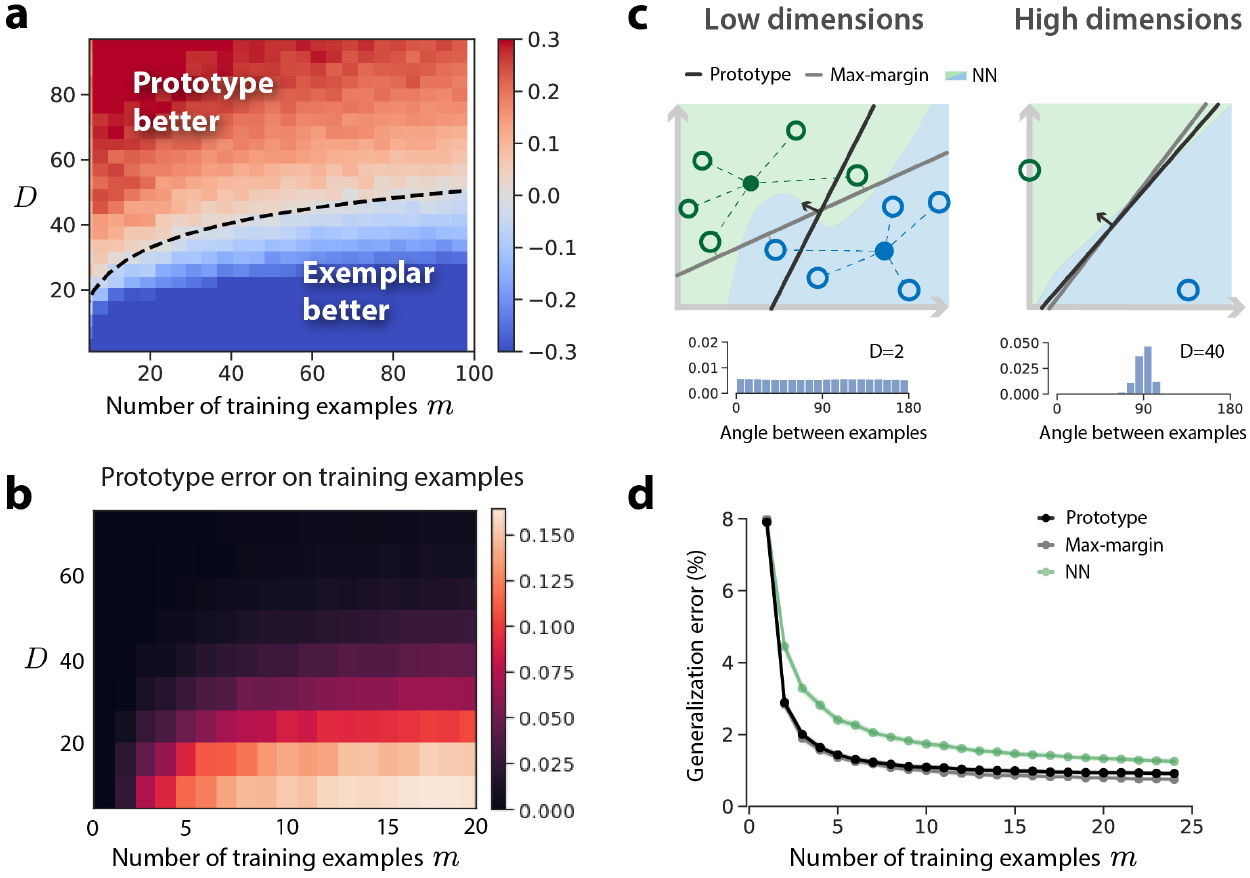
Comparing cognitive learning models on naturalistic tasks. **a**, We compare the performance of prototype and exemplar learning as a joint function of concept manifold dimensionality and the number of training examples, using novel visual concept manifolds from a trained ResNet50. We vary dimensionality by projecting concept manifolds onto their top *D* principal components. We formalize exemplar learning by a nearest-neighbors decision rule (NN, see SI 5). Because the generalization error is very close to zero for *m* and *D* large, here we plot SNR^proto^ − SNR^NN^. The dashed line demarcating the boundary is given by *m* = exp(*D/*10), reflecting the relationship log *m* ∝ *D* predicted by our theory (SI 5). The constant 10 is chosen by a one-parameter fit. **b**, Prototype learning error as a function of *m* and *D* when training examples are re-used as test examples. A prototype classifier may misclassify one or more of the training examples themselves when concepts are low dimensional and the number of training examples is large. However this rarely happens in high dimensions, as as illustrated in **c. c**, In low dimensions, multiple training examples may overlap along the same direction (inset histogram; distribution of angles between examples in DNN concept manifolds for *D* = 2). Hence averaging these examples (open circles) to yield two prototypes (solid circles) may leave some of the training examples on the wrong side of the prototype classifier’s decision boundary (black line). In high dimensions, however, all training examples are approximately orthogonal (inset histogram, *D* = 40), so such mistakes rarely happen. Panel **c** also shows the decision boundaries of max-margin learning (grey line) and NN learning (green and blue regions), both of which perfectly classify the training examples. In low dimensions prototype and max-margin learning may learn very different decision boundaries; however in high dimensions their decision boundaries are very similar, as quantified in **d. d**, Empirical comparison of prototype, max-margin, and NN exemplar learning as a function of the number of training examples *m* (Methods 3.8). When the number of training examples is small, prototype and SVM learning are approximately equivalent. For larger *m*, SVM outperforms prototype learning. NN learning performs worse than SVM and prototype learning for both small and intermediate *m*.

An intermediate model between prototype and exemplar learning is max-margin learning^42^ (Fig. 8**c**). Like prototype learning, max-margin learning involves learning a linear readout; however, rather than pointing between the concept prototypes, its linear readout is chosen to maximize the distance from the decision boundary to the nearest training example of each concept^42^ (Supp. Fig. 8). Max-margin learning is more sophisticated than prototype learning in that it incorporates not only the estimated manifold centroid but also the variation around the centroid. Thus it is able to achieve zero generalization error for large *m* when concept manifolds are linearly separable, overcoming the limitation on the prototype learning SNR due to signal-noise overlaps in the large-*m* limit (Eq. 1, Fig. 2**d**). However, like exemplar learning it requires memory of all training examples. Comparing the three learning models on DNN concept manifolds we find that prototype and max-margin learning are approximately equivalent for *m* ≲ 8, and both outperform exemplar learning for *m* of small to moderate size (Fig. 8**d**, Methods 3.8).

### 2.9 Discussion

We have developed a theoretical framework that accounts for the remarkable accuracy of human few-shot learning of novel high-dimensional naturalistic concepts in terms of a simple, biologically plausible neural model of concept learning. Our framework defines the concepts that we can rapidly learn in terms of tight manifolds of neural activity patterns in a higher-order brain area. The theory provides new, readily measurable geometric properties of population responses (Fig. 2) and successfully links them to the few-shot learning performance of a simple prototype learning model. Our model yields remarkably high accuracy on few-shot learning of novel naturalistic concepts using deep neural network representations for vision (Fig. 1 and Fig. 3) which is in excellent quantitative agreement with theoretical prediction. We further show that the four geometric quantities identified by our theory undergo orchestrated transformations along the layers of trained DNNs, and along the macaque ventral visual pathway, yielding a consistent improvement in performance along the system’s hierarchy (Fig. 4 and Fig. 5). We extend our theory to cross-domain learning, and demonstrate that comparably powerful visual concept learning is attainable from linguistic descriptors of concepts *without* visual examples, using a simple map between language and visual domains (Fig. 7). We show analytically and confirm numerically that high few-shot learning performance is possible with as few as 200 IT-like neurons (Fig. 6).

#### A design tradeoff governing neural dimensionality

A surprising result of our theory is that rapid concept learning is easier when the underlying variability of images belonging to the same object is spread across a large number of dimensions (Fig. 2**c**,**g**). Thus, high-dimensional neural representations allow new concepts to be learned from fewer examples, while, as has been shown recently, low-dimensional representations allow for a greater number of familiar concepts to be classified^30,31^. Understanding the theoretical principles by which neural circuits negotiate this tradeoff constitutes an important direction for future research.

#### Asymmetry in few-shot learning

We found that the pairwise generalization error of simple cognitive models like prototype, exemplar, and max-margin learning exhibits a dramatic asymmetry when only one or a few training examples are available (Supp. Fig. 1). Our theory attributes this asymmetry to a bias arising from the inability of simple classifiers to estimate the variability of novel concepts from a few examples (Fig. 2**b**,**f**). Investigating whether humans exhibit the same asymmetry, or identifying mechanisms by which this bias could be overcome, are important future directions. An interesting hypothesis is that humans construct a prior over the variability of novel concepts, based on the variability of previously learned concepts^43^.

#### Connecting language and vision

Remarkably, we found that machine learning derived language representations and visual representations, despite being optimized for different tasks across different modalities, can be linked together by an exceedingly simple linear map (Fig. 5), to enable learning novel visual concepts given only language descriptions. Recent work has revealed that machine learning derived neural language representations match those in humans as measured by both ECOG and fMRI^44^. Our results suggest that language and high-level visual representations of concepts in humans may indeed be related through an exceedingly simple map, a prediction that can be tested experimentally. Broadly, our results add plausibility to the tenets of dual-coding theory in human cognition^45^.

#### Comparing brains and machines

Computational neuroscience has recently developed increasingly complex high-dimensional machine learning-derived models of many brain regions, including the retina^46–49^, V1, V4 and IT^22,50^, motor cortex^51^, prefrontal cortex^52^, and enthorhinal cortex^53,54^. Such increased model complexity raises foundational questions about the appropriate comparisons between brains and machine based models^55^. Previous approaches based on behavioral performance^16,25,56–58^, neuron^46^ or circuit^49^ matching, linear regression between representations^22^, or representational similarity analysis^24^, reveal a reasonable match between the two. However, our higher-resolution decomposition of performance into a fundamental set of observable geometric properties reveals significant mismatches (Fig. 4 and Fig. 6**d**). In particular, intermediate representations corresponding to V4 have much lower dimension and higher signal in macaque compared to DNNs, calling for more veridical models of the visual pathway and a better understanding of visual processing in V4.

#### Comparing cognitive learning models on naturalistic concepts

Our theory reveals that exemplar learning is superior to prototype learning given many examples of low-dimensional concepts, consistent with past laboratory experiments^12,39,40,59^, but is inferior to prototype learning given only a few examples of high-dimensional concepts, like those in DNNs and in primate IT (Fig. 8), shedding light on a 40-year-old debate^5^. These predictions are consistent with a recent demonstration that a prototype-based rule can match the performance of an exemplar model on categorization of familiar high dimensional stimuli^58^. We go beyond prior work by (1) demonstrating that prototype learning achieves superior performance on few-shot learning of novel naturalistic concepts, (2) precisely characterizing the tradeoff as a joint function of concept manifold dimensionality and the number of training examples (Fig. 8), and (3) offering a theoretical explanation of this behavior in terms of the geometry of concept manifolds (SI 5).

#### Proposals for experimental tests of our model

Our theory makes specific predictions that can be tested through behavioral experiments designed to evaluate human performance at learning novel visual concepts from few examples (see Supp. Fig. 7 for a proposed experimental design). First, we predict a specific pattern of errors across these novel concepts, shared by neural representations in several trained DNNs (proxies for human IT cortex) as well as neural representations in macaque IT (Fig. 5 and Supp. Fig. 7**c**). Second, we predict that humans will exhibit a marked asymmetry in pairwise few-shot learning performance following the pattern derived in our theory (see Supp. Fig. 7 for examples). Third, we predict how performance should scale with the number of training examples *m* (Fig. 8**d**). Matches between these predictions and experimental results would indicate that simple classifiers learned atop IT-like representations may be sufficient to account for human concept learning performance. Deviations from these predictions may suggest that humans leverage more sophisticated higher-order processing to learn new concepts, which may incorporate human biases and priors on object-like concepts^43^.

By providing new fundamental links between the geometry of concept manifolds in the brain and the performance of few-shot concept learning, our theory lays the foundations for next generation combined physiology and psychophysics experiments. Simultaneously recording neural activity and measuring behavior would allow us to test our hypothesis that the proposed neural mechanism is sufficient to explain few-shot learning performance, and to test whether the four fundamental geometric quantities we identify correctly govern this performance. Furthermore, ECOG or fMRI could be used to investigate whether these four geometric quantities are invariant across primates and humans. Conceivably, our theory could even be used to design visual stimuli to *infer* the geometry of neural representations in primates and humans, without the need for neural recordings.

In conclusion, this work represents a significant step towards understanding the neural basis of concept learning in humans, and proposes theoretically guided psychophysics and physiology experiments to further illuminate the remarkable human capacity to learn new naturalistic concepts from few examples.

## 3 Methods

### 3.1 Visual stimuli

Visual stimuli were selected from the ImageNet dataset, which includes 21, 840 unique visual concepts. A subset of 1, 000 concepts comprises the standard ImageNet1k training set used in the ILSVRC challenge^60^. All DNNs studied throughout this work are trained on these 1, 000 concepts alone. To evaluate few-shot learning on novel visual concepts, we gather an evaluation set of 1, 000 concepts from the remaining 20, 840 concepts not included in the training set, as follows. The ImageNet dataset is organized hierarchically into a semantic tree structure, with each visual concept a node in this tree. We include only the leaves of the tree (e.g. ‘wildebeest’) in our evaluation set, excluding all superordinate categories (e.g. ‘mammal’) which have many descendants. We additionally exclude leaves which correspond to abstract concepts such as ‘green’ and ‘pet’. Finally, the concepts in the ImageNet dataset vary widely in the number of examples they contain, with some concepts containing as many as 1, 500 examples, and others containing only a single example. Among the concepts that meet our criteria above, we select the 1, 000 concepts with the greatest number of examples. A full list of the 1, 000 concepts used in our evaluation set is available at github.com/bsorsch/geometry_fewshot_learning.

### 3.2 Estimating the geometry of concept manifolds

Each visual stimulus elicits a pattern of activity ***x*** ∈ ℝ^*N*^ across the sensory neurons in a high-order sensory layer such as IT cortex, or analogously the feature layer of a DNN. We collect the population responses to each of all *P* images in the dataset belonging to a particular visual concept in an *N* × *P* matrix *X*. To estimate the geometry of the underlying concept manifold, we construct the empirical covariance matrix 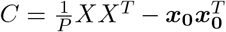, where ***x***_**0**_ ∈ ℝ^*N*^ is the manifold centroid. We then diagonalize the covariance matrix,

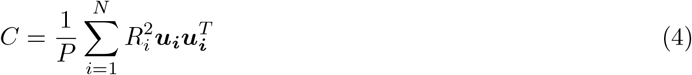

Where ***u***_***i***_ are the eigenvectors of *C* and 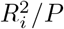 the associated eigenvalues. The ***u***_*i*_ each represent unique, potentially interpretable visual features (e.g. animate vs inanimate, spiky vs stubby, or short-haired vs long-haired^19^). Individual examples of the concept vary from the average along each of these ‘noise’ directions. Some noise directions exhibit more variation than others, as governed by the *R*_*i*_. geometrically, the eigenvectors ***u***_***i***_ correspond to the principal axes of a high-dimensional ellipsoid centered at ***x***_**0**_, and the *R*_*i*_ correspond to the radii along each axis. A useful measure of the total variation around the centroid is the mean squared radius, 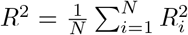, *R*^2^ sets a natural length scale, which we use in our theory as a normalization constant to obtain interpretable, dimensionless quantities. Although we do not restrict the maximal number of available axes, which could be as large as the dimensionality of the ambient space, *N*, the number of directions along which there is significant variation is quantified by an effective dimensionality 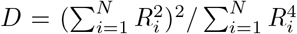, called the participation ratio^29^, which in practical situations is much smaller than *N*. The participation ratio arises as a key quantity in our theory (SI 2.3).

### 3.3 Prototype learning

To simulate *m*-shot learning of novel concepts pairs, we present *m* randomly selected training examples of each concept to our model of the visual pathway, and collect their neural representations ***x***^*aµ*^, ***x***^*bµ*^, *µ* = 1, …, *m* in a population of IT-like neurons. We perform prototype learning by averaging the representations of each concept into concept prototypes, 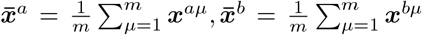. To evaluate the generalization accuracy, we present a randomly selected test example of concept *a*, and determine whether its neural representation is closer in Euclidean distance to the correct prototype 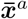 than it is to the incorrect prototype 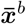. This classification rule can be implemented by a single downstream neuron which adjusts its synaptic weight vector ***w*** to point along the difference between the two concept prototypes, 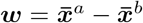, and adjusts its bias (or firing rate threshold) *β* to equal the average overlap of ***w*** with each prototype, 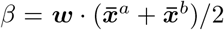. We derive an analytical theory for the generalization error of prototype learning on concept manifolds in SI 2.3, and we extend our model and theory to classification tasks involving more than two concepts in SI 2.4.

### 3.4 Prototype learning experiments on synthetic concept manifolds

To evaluate our geometric theory, we perform prototype learning experiments on synthetic concept manifolds constructed with pre-determined ellipsoidal geometry (Fig. 2**e-h**). By default, we construct manifolds with 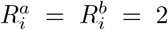, *i* = 1,…,*D* and *D* = 50, and we sample the centroids 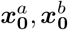 and subspaces 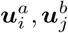 randomly under the constraint that the signal direction **Δ*x***_**0**_ and the subspace directions are orthonormal,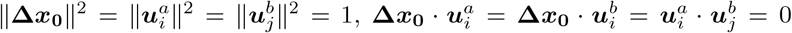, so that the signal-noise overlaps are zero. We then vary each geometric quantity individually, over the ranges reflected in Fig. 2**e-h**. To vary the signal, we vary 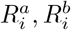 from 1 to 2.5. To vary the bias over the interval (−1, 1), we fix 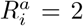 and vary 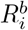 from 0 to 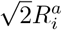. To vary the signal-noise overlaps, we construct ellipsoidal manifolds with one long direction, 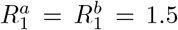, and *D* − 1 short directions, 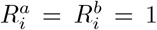, *i* = 2,…,*D*. We then vary the angle *θ* between the signal direction **Δ*x***_**0**_ and the first subspace basis vector 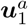 by choosing 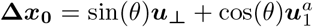, where 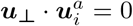. We vary *θ* from fully orthogonal (*θ* = *π/*2) to fully overlapping (*θ* = 0).

### 3.5 DNN concept manifolds

All DNNs studied throughout this work are standard architectures available in the PyTorch library^61^, and are pretrained on the ImageNet1k dataset. To obtain novel visual concept manifolds, we randomly select *P* = 500 images from each of the *a* = 1, …, 1000 never-before-seen visual concepts in our evaluation set, pass them into the DNN, and obtain their representations in the feature layer (final hidden layer). We collect these representations in an *N* × *P* response matrix *X*^*a*^. *N* = 2048 for ResNet architectures. For architectures with *M* > 2048 neurons in the feature layer, we randomly project the representations down to *N* = 2048 dimensions using a random matrix *A* ∈ ℝ^*N ×M*^, 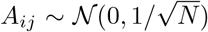. We then compute concept manifold geometry as described in Methods 3.2. To study the layerwise behavior of manifold geometry, we collect the representations at each layer *l* of the trained DNN into an *N* × *P* matrix 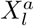. For the pixel layer, we unravel raw images into 224 224 3 dimensional vectors and randomly project down to *N* dimensions. Code is available at github.com/bsorsch/geometry_fewshot_learning.

### 3.6 Macaque neural recordings

Neural recordings of macaque V4 and IT were obtained from the dataset collected in Majaj et al.^21^. This dataset contains 168 multiunit recordings in IT and 88 multiunit recordings in V4 in response to 3,200 unique visual stimuli, over ∼ 50 presentations of each stimulus. Each visual stimulus is an image of one of 64 distinct synthetic 3d objects, randomly rotated, positioned, and scaled atop a random naturalistic background. To obtain the concept manifold for object *a*, we collect the average response of IT neurons to each of the *P* = 50 unique images of object *a* in an *N*_IT_ × *P* response matrix 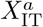, where *N*_IT_ = 168, and compute the geometry of the underlying manifold as described in Methods 3.2. We repeat for V4, obtaining a *N*_V4_ × *P* response matrix *X*_V4_, where *N*_V4_ = 88. We additionally simulate V1 neural responses to the same visual stimuli via a biologically constrained Gabor filter bank, as described in Dapello et al.^62^ To compare with trained DNNs, we pass the same set of stimuli into each DNN, and obtain a *N*_DNN_ *× P* response matrix *X*_DNN_, as described in Methods 3.5. We then project this response matrix into *N*_IT_, *N*_V4_, or *N*_V1_ dimensions to compare with IT, V4, or V1. In order to align V1,V4, and IT to corresponding layers in the trained DNN (as in Fig. 5**d**), we identify the DNN layers that are most predictive of V1, V4, and IT using partial least squares regression with 25 components, as in Schrimpf et al.^28^

### 3.7 Visual concept learning without visual examples by aligning visual and language domains

To obtain language representations for each of the 1,000 familiar visual concepts from the ImageNet1k training dataset, we collected their embeddings in a standard pre-trained word vector embedding model (GloVe)^36^. The word vector embedding model is pre-trained to produce neural representations for each word in the English language based on word co-occurrence statistics. Since concept names in the ImageNet dataset typically involve multiple words (e.g. “tamandua, tamandu, lesser anteater, Tamandua tetradactyla”) we averaged the representations for each word in the class name into a single representation 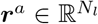, where *N*_*l*_ = 300 (see Supp. Fig. 4 for an investigation of this choice). We collected the corresponding visual concept manifolds in a pre-trained ResNet50. To align the language and vision domains, we gathered the centroids of the *a* = 1, …, 1, 000 training manifolds 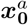 into an *N ×* 1, 000 matrix *X*_0_, and gathered the corresponding language representations into an *N*_*L*_ × 1, 000 matrix *Y*. We then learned a scaled linear isometry *f* from the language domain to the vision domain by solving the generalized Procrustes analysis problem *f* = min_*α,O*,***b***_ ‖*f*_*α,O*,***b***_(*Y*) − *X*_0_‖ ^2^, where *f*_*α,O*,***b***_ = *αOY* − ***b***, *α* is a scalar, 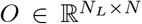 is an orthogonal matrix, and ***b*** ∈ ℝ^*N*^ is a translation.

### 3.8 Comparing cognitive learning models on naturalistic tasks

In addition to prototype learning, we performed few-shot learning experiments using two other decision rules: a max-margin classifier and a nearest-neighbors classifier (Fig. 8). The max-margin classifier was optimized using a standard support vector machine (SVM) software package, LibSVM^63^, using a linear kernel and an *L*_2_ regularization constant *C* = 5 × 10^4^. The nearest-neighbors classifier was evaluated by computing the Euclidean distance of a test example to each training example, and categorizing the test example according to the identity of the nearest training example. Exemplar learning more generally allows for comparisons to more than just the nearest neighbor, and involves the choice of a parameter *β* which weights the contribution of each training example to the discrimination function. When *β* =∞, only the nearest neighbor contributes to the discrimination function. When *β* = 0, all training examples contribute equally. However, we find that the nearest-neighbors limit *β* → ∞ is close to optimal in our setting (Fig. 8). Hence we formalize exemplar learning by a nearest-neighbors decision rule. In order to study the effect of concept manifold dimensionality *D* on the performance of each learning rule (Fig. 8**a**,**b**), we vary *D* by projecting concept manifolds onto their top *D* principal components.

## 4 Acknowledgements

The work is partially supported by the Gatsby Charitable Foundation, the Swartz foundation, and the National Institutes of Health (Grant No. 1U19NS104653). B.S. thanks the Stanford Graduate Fellowship for financial support. S.G. thanks the Simons and James S McDonnell foundations, and an NSF CAREER award.

## 5 Supplemental figures

**Supplementary Figure 1:**
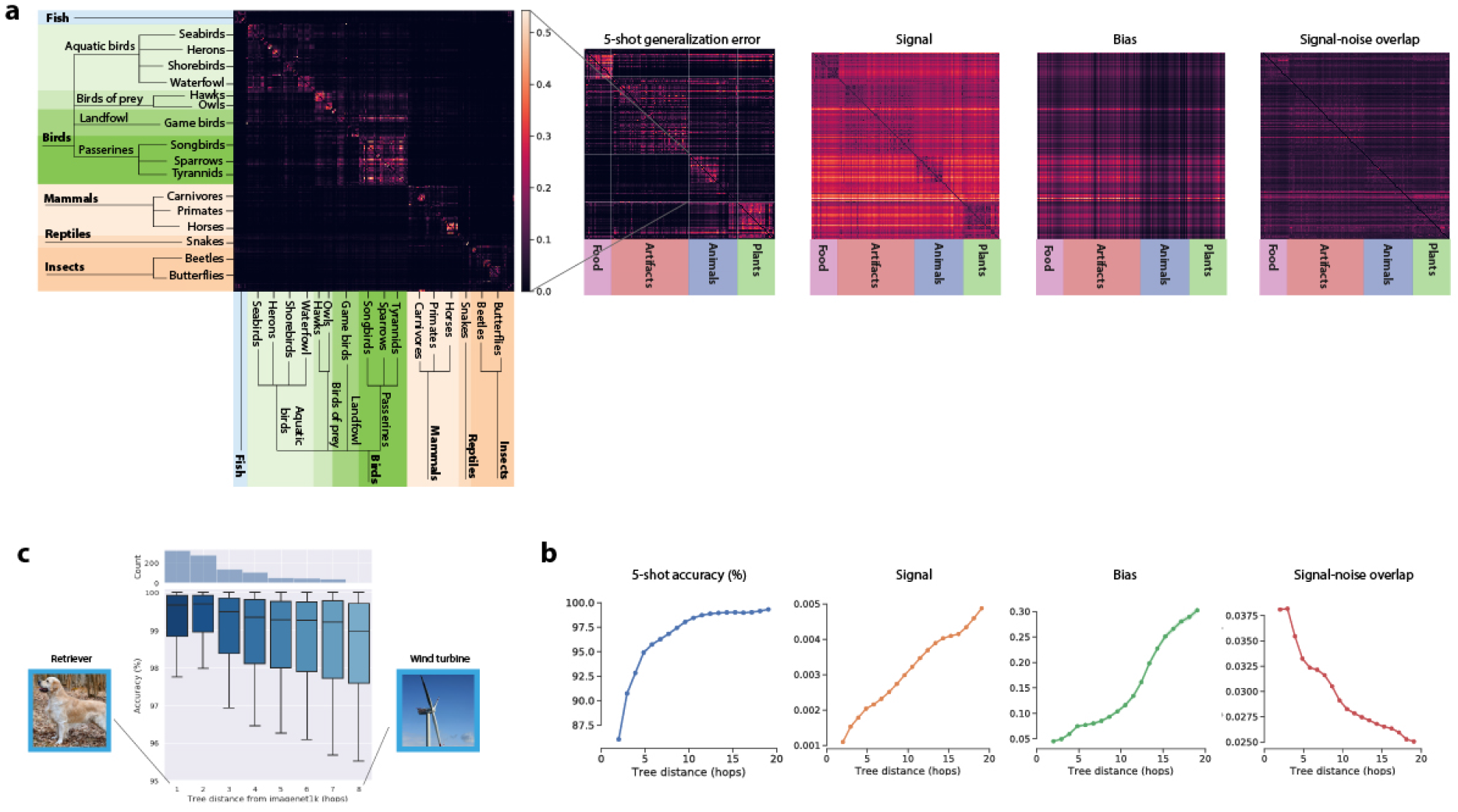
Geometry of DNN concept manifolds encodes a rich semantic structure. See SI 6. **a**, We sort the generalization error pattern of prototype learning using concept manifolds from a trained ResNet50 to obey the hierarchical semantic structure of the ImageNet21k dataset. The sorted error matrix exhibits a prominent block diagonal structure, suggesting that most of the errors occur between concepts on the same branch of the semantic tree, and errors between two different branches of the semantic tree are exceedingly unlikely. *Inset:* error pattern across a subset of novel visual concepts, including fish birds, mammals, reptiles and insects. The full error pattern across all 1,000 novel visual concepts is shown at right. Rows correspond to concepts from which test examples are drawn. This error pattern exhibits a pronounced asymmetry, with much larger errors above the diagonal than below. For instance, food and artifacts are more likely to be classified as plants or animals than plants and animals are to be classified as food or artifacts. We additionally plot the sorted pattern of individual geometric quantities: signal, bias, and signal-noise overlap. Signal exhibits a pronounced block diagonal structure, similar to the error pattern. Bias exhibits a pronounced asymmetry, indicating that plant and animal concept manifolds have significantly smaller radii than artifact and food concept manifolds do. **b**, We plot the average few-shot accuracy, signal, bias, and signal-noise overlap across all pairs of concepts, as a function of the distance between the two concepts on the semantic tree, defined as the number of hops required to travel from one concept to the other. Few-shot learning accuracy, signal, and bias all increase significantly with semantic distance, while signal-noise overlaps decrease. **c**, To quantify the effect of distribution shift from the training concepts to the novel concepts, we measure the tree distance from each of the 1k novel concepts to its nearest neighbor among the 1k training concepts. We plot the average few-shot learning accuracy as a function of this distance. Few-shot learning accuracy degrades slightly with distance from the training set, but the effect is not dramatic.

**Supplementary Figure 2:**
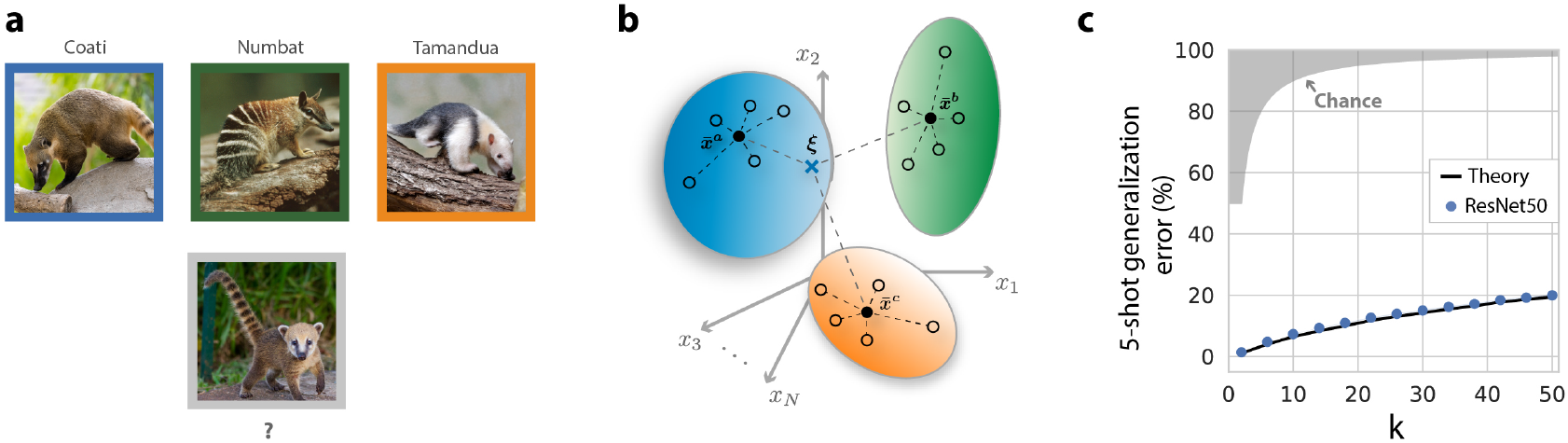
Learning many novel concepts from few examples. Concept learning often involves categorizing more than two novel concepts. In SI 2.4 we extend our theory to model few-shot learning of *k* novel concepts. **a**, An example one-shot learning task for *k* = 3: does the test image in the gray box contain a ‘coati’ (blue box), a ‘numbat’ (green box), or a ‘tamandua’ (orange box), given one training example of each? **b**, Illustration of *k*-concept learning. Training examples of each novel concept (open circles) are averaged into *k* class prototypes (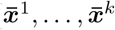 solid circles). A test example (***ξ***, blue cross) is classified based on its Euclidean distance to each of the concept prototypes. This classification can be performed by *k* downstream neurons, one for each novel concept, which adjust their synaptic weights to point along the concept prototypes. **b**, Empirical performance and theoretical predictions. We perform 5-shot learning experiments on visual concept manifolds extracted from a DNN in response to 1, 000 novel visual concepts from the ImageNet21k dataset. We compute the generalization error as a function of the number of novel concepts to be learned, *k*, as well as the prediction from our theory (SI 2.4). Performance is remarkably high, and generalization error stays below 20% even for *k* = 50 (where error at chance is 98%).

**Supplementary Figure 3:**
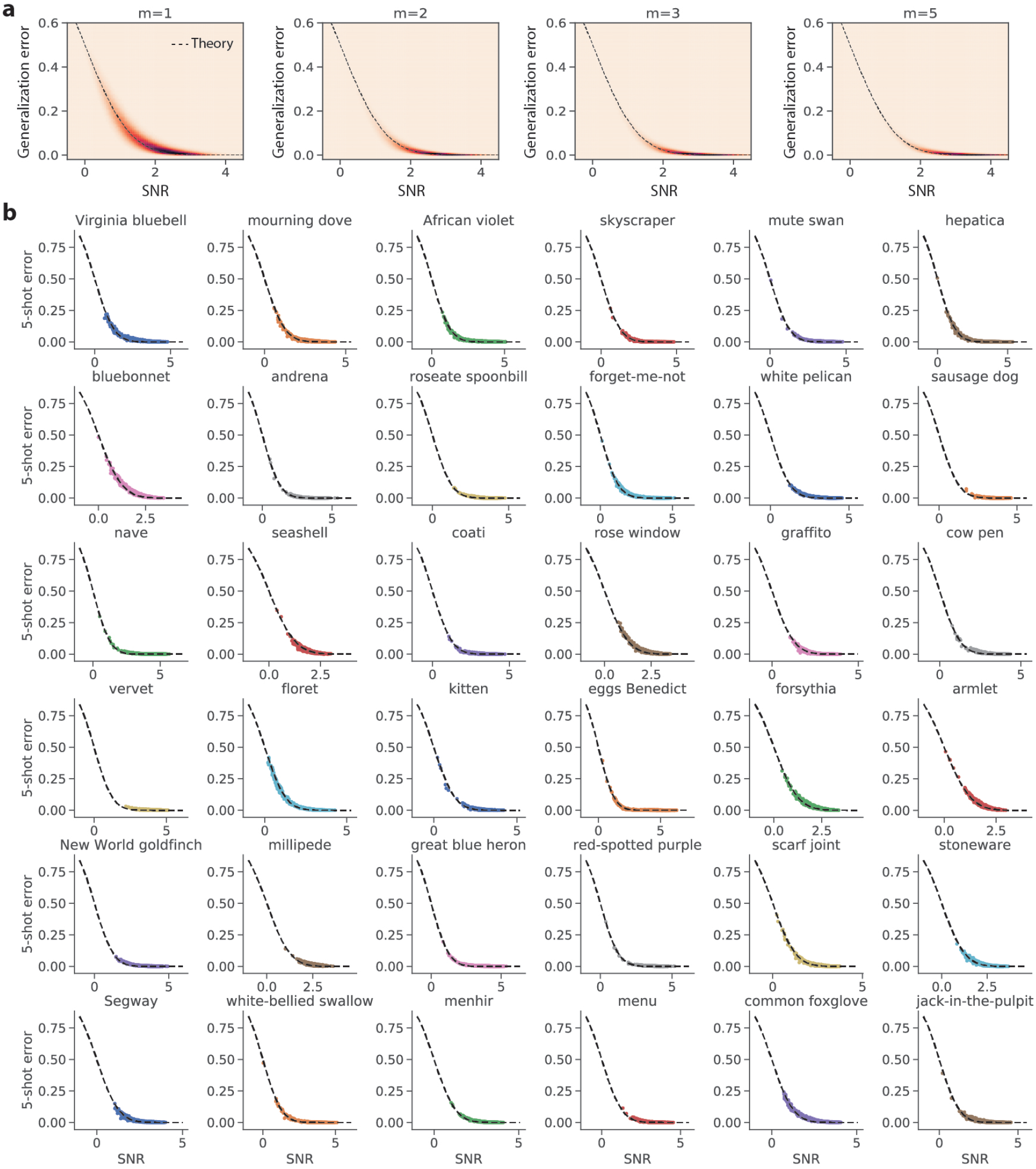
geometric theory and few-shot learning experiments on a variety of novel concepts. **a**, We compare the empirical generalization error in 1–, 2–, 3–, and 5-shot learning experiments to the prediction from our geometric theory (Eq. SI.38) on all 1, 000 × 999 pairs of visual concepts from the ImageNet21k datset, using concept manifolds derived from a trained ResNet50. We plot a 2d histogram rather than a scatterplot because the number of points is so large. *x-axis*: SNR obtained by estimating neural manifold geometry. *y-axis*: Empirical generalization error measured in few-shot learning experiments. Theoretical prediction (dashed line) shows a good match with experiments. **b**, We provide additional examples of 5-shot prototype learning experiments in a ResNet50 (colored points), along with the prediction from our geometric theory (dashed line), on 36 randomly selected novel visual concepts from the ImageNet21k dataset. Each panel plots the generalization error of one novel visual concept (e.g. ‘Virginia bluebell’) against all 999 other novel visual concepts (e.g. ‘bluebonnet’, ‘African violet’). Each point represents the average generalization error on one such pair of concepts. *x-axis*: SNR (Eq. 1) obtained by estimating neural manifold geometry. *y-axis*: Empirical generalization error measured in few-shot learning experiments. Theoretical prediction (dashed line) shows a good match with experiments. Error bars, computed over many draws of the training and test examples, are smaller than the symbol size..

**Supplementary Figure 4:**
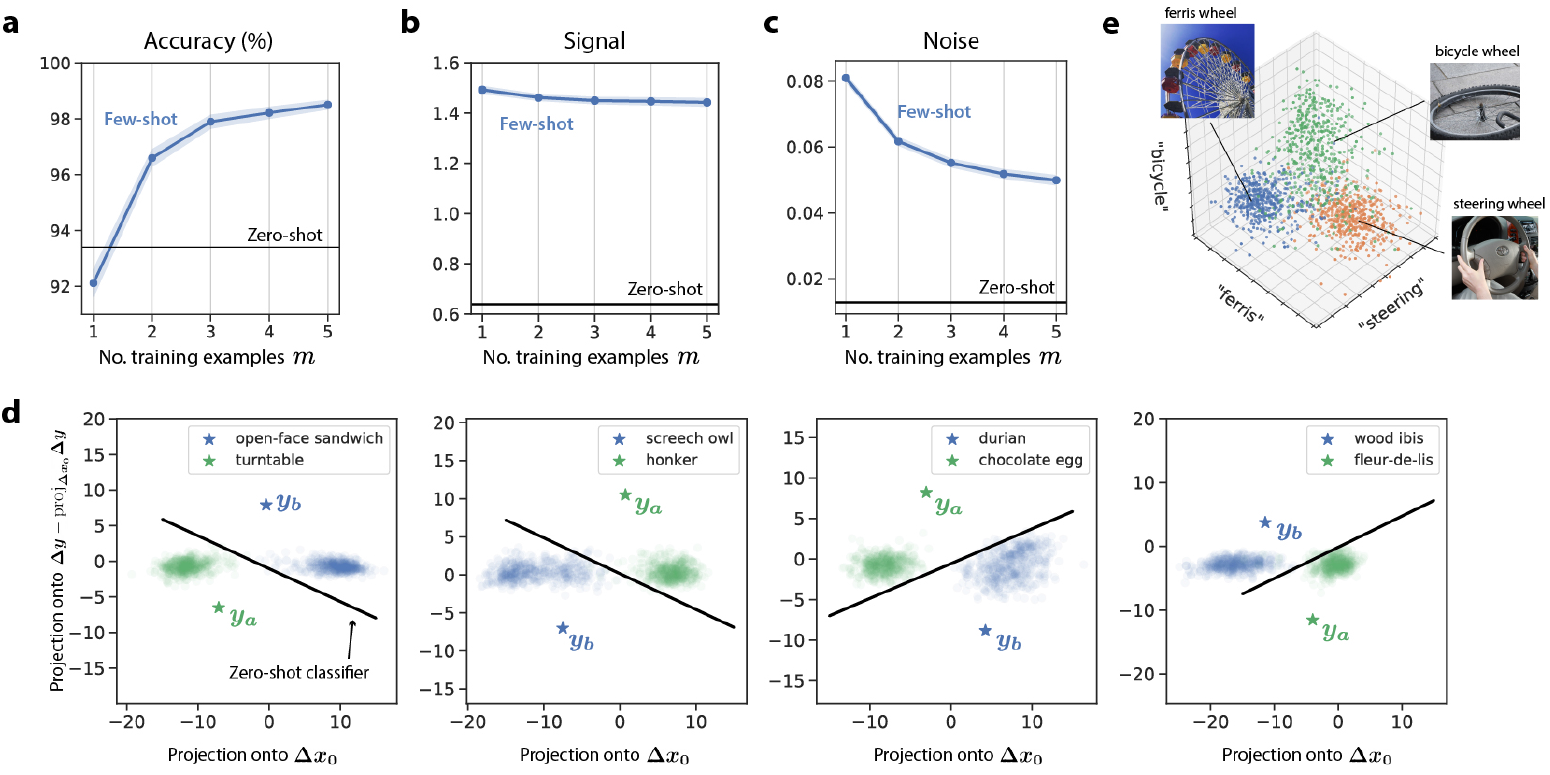
How many words is a picture worth? Comparing prototypes derived from language and vision. See SI 3.2. **a**, We compare the performance of prototype learning using prototypes derived from language representations (*zero-shot learning*, Sec. 2.7) to those derived from one or a few visual examples (*few-shot learing*, Sec. 2.1). We find that prototypes derived from language yield a better generalization accuracy than those derived from a single visual example, but not two or more visual examples. **b**,**c**,**d**, To better understand this behavior, we use our geometric theory for zero-shot learning, Eq. 3, to decompose the performance of zero- and few-shot learning into a contribution from the ‘signal’, which quantifies how closely the estimated prototypes match the true concept centroids, and a contribution from the ‘noise’, which quantifies the overlap between the readout direction and the noise directions. We find that both signal, **b**, and noise, **c**, are significantly lower for zero-shot learning than for few-shot learning. Hence one-shot learning prototypes more closely match the true concept prototypes on average than language prototypes do. However, language prototypes are able to achieve a higher generalization accuracy by picking out readout directions which overlap significantly less with the concept manifolds’ noise directions. **d**, To visualize this, we project pairs of concept manifolds into the two-dimensional space spanned by the difference between the manifold centroids, **Δ*x***_**0**_, and the language prototype readout direction, **Δ*y***. Blue and green stars indicate the language-derived prototypes, and the black boundary indicates the zero-shot learning classifier which points between the two language prototypes. Each panel shows a randomly selected pair of concepts. In each case, the manifolds’ variability is predominantly along the **Δ*x***_**0**_ direction, while the language prototypes pick out readout directions **Δ*y*** with much lower variability. **e**, To obtain a single language representation for visual concepts with multiple word labels (e.g. ‘ferris wheel’, ‘bicycle wheel’, ‘steering wheel’), we chose to simply average the representations of each word. This choice only succeeds if the modifying words (e.g. ‘ferris’, ‘bicycle’, ‘steering’) correspond to meaningful directions when mapped into the visual representation space. We investigate this choice visually by projecting the ‘ferris wheel’, ‘bicycle wheel’, and ‘steering wheel’ visual concept manifolds into the three-dimensional space spanned by the word representations for ‘ferris’, ‘bicycle’, and ‘steering’ mapped into the visual representation space. We find that the three concept manifolds are largely linearly discriminable in this three-dimensional space, indicating that averaging the word representations can be an effective strategy, though likely not the optimal choice.

**Supplementary Figure 5:**
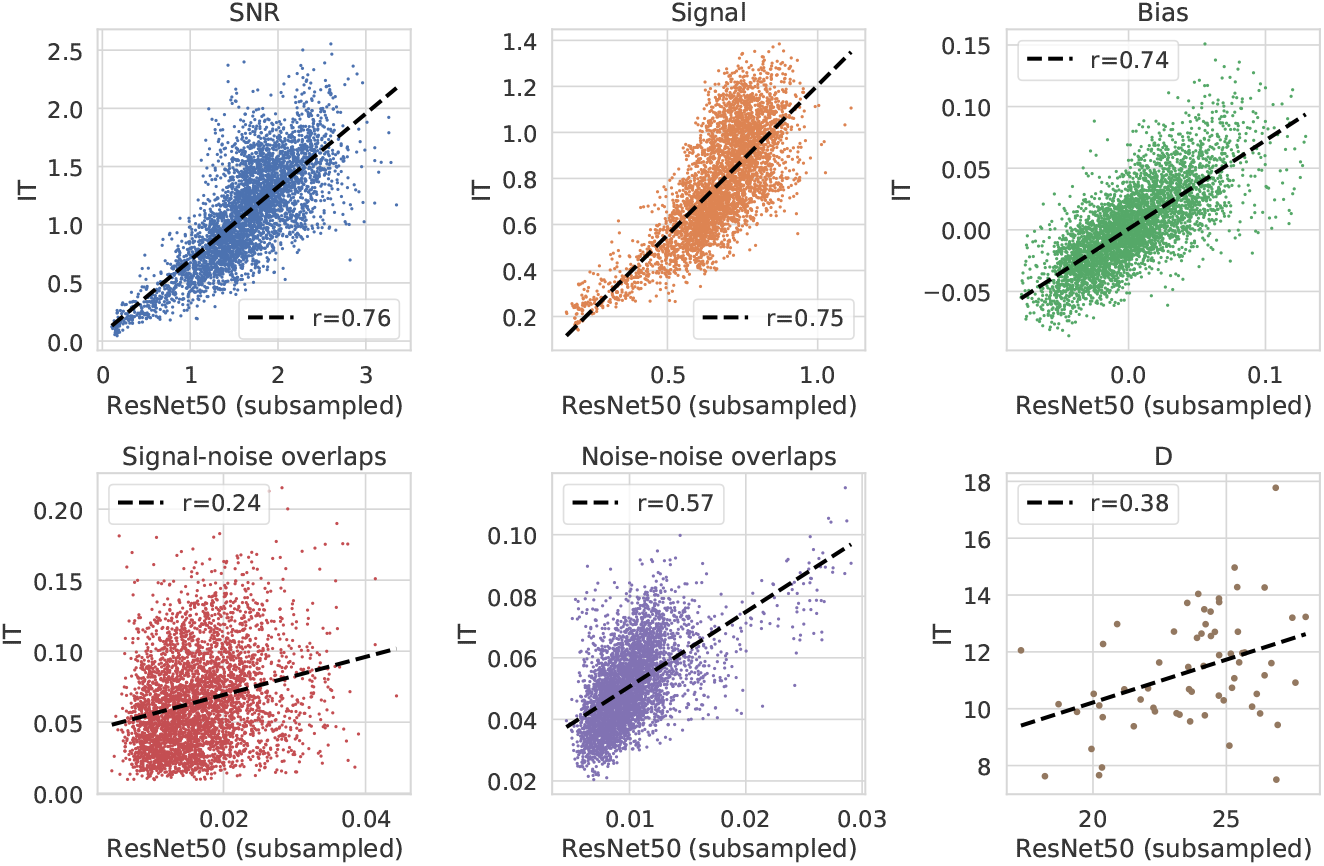
Concept manifold geometry is correlated across primate IT cortex and trained DNNs. We estimate the geometry of visual concept manifolds in primate IT cortex and in trained DNNs in response to the same 64 naturalistic visual concepts^21^. We then compute the correlation between each quantity in IT cortex and in a trained DNN. Here we use a ResNet50, whose neurons have been randomly subsampled to match the number of recorded neurons in macaque IT (168 neurons). Each panel shows one geometric quantity: SNR (r=0.76, *p <* 1 × 10^−10^), signal (r=0.75, *p <* 1 × 10^−10^), bias (r=0.74, *p <* 1 × 10^−10^), signal-noise overlaps (r=0.24, *p <* 1 × 10^−10^), noise-noise overlaps (see SI 2.3; r=0.57, *p <* 1 × 10^−10^), and dimension (*r* = 0.38, *p <* 0.005).

**Supplementary Figure 6:**
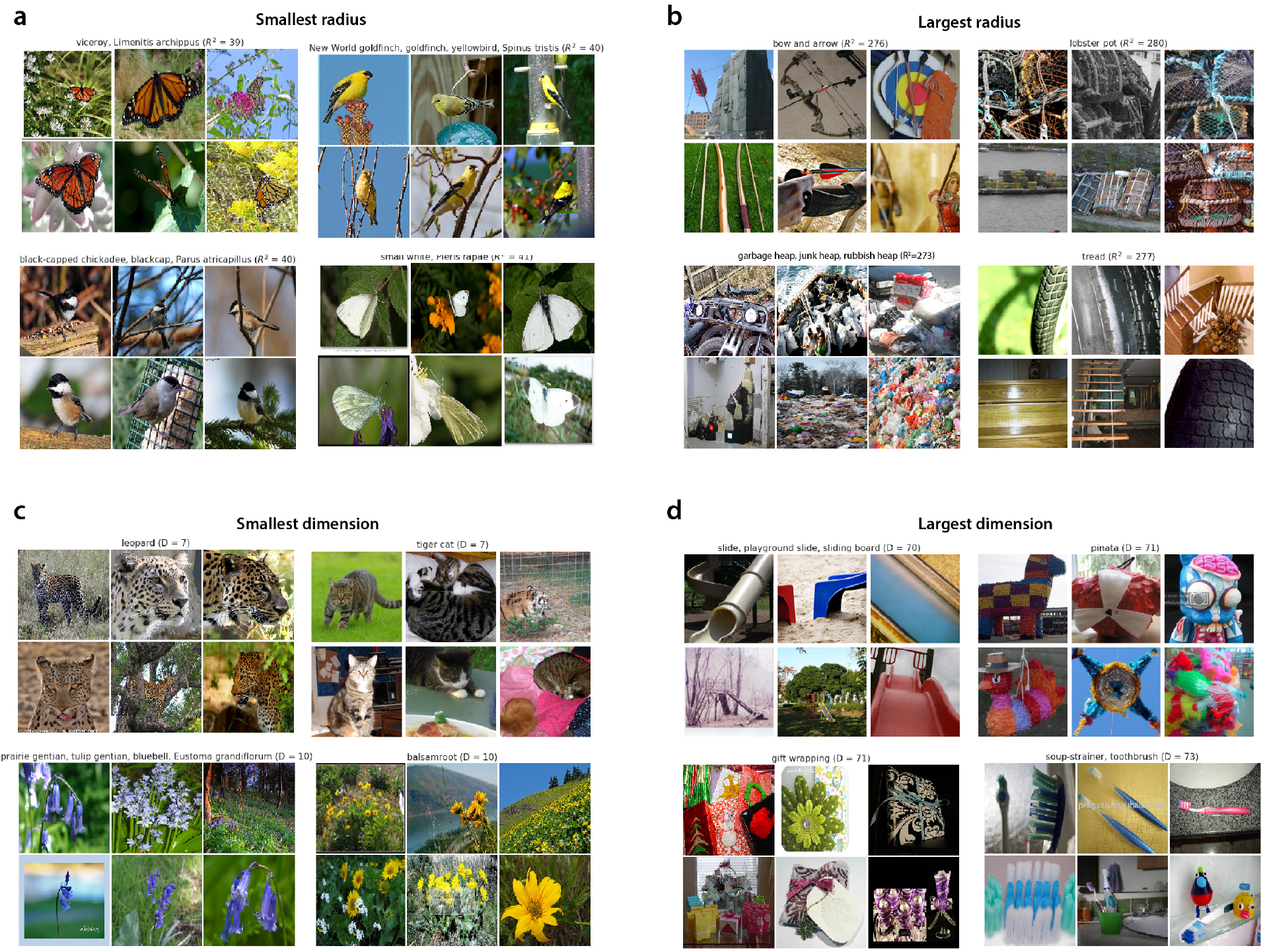
Visual examples of concept manifolds with small and large dimension and radius. Among the 1, 000 novel visual concepts in our heldout set, we collect examples of the visual concepts whose manifolds in a trained ResNet50 have, **a**, smallest radius, **b**, largest radius, **c**, smallest dimension, and **d**, largest dimension. The salient visual features of concepts with small manifold radius, **a**, appear to exhibit significantly less variation than those of concepts with large manifold radius, **b**. Furthermore, we observe that the visual concepts with smallest manifold radius and dimension are largely comprised of plants and animals **a**,**c**, while the visual concepts with largest manifold and dimension are largely comprised of human-made objects **b**,**d**.

**Supplementary Figure 7:**
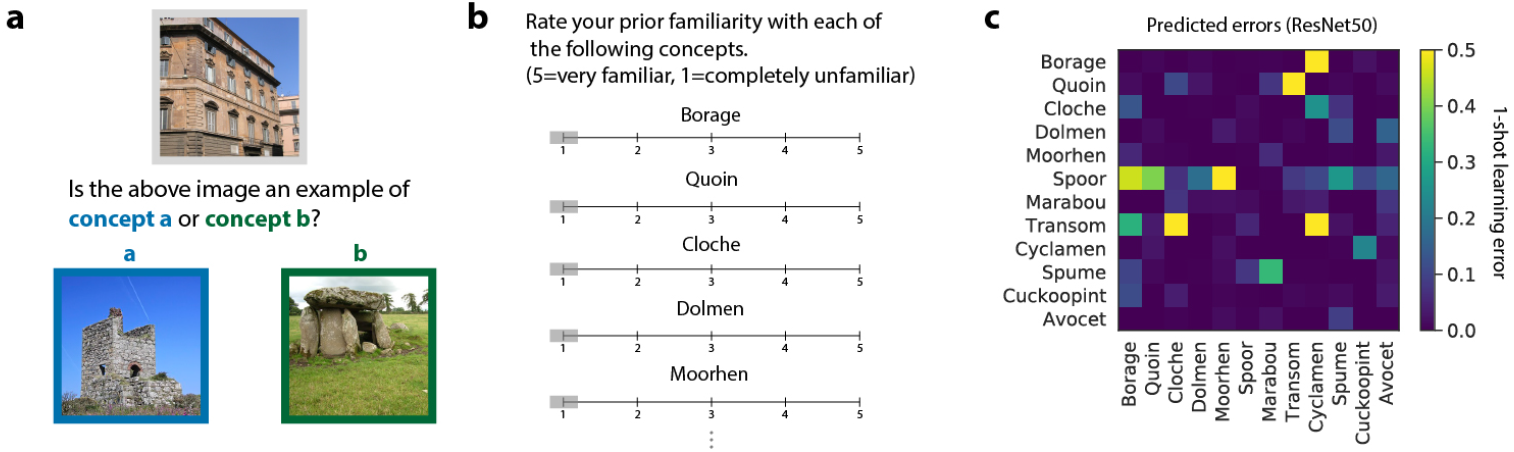
Proposed psychophysics experiment to evaluate human few-shot learning on novel naturalistic concepts.. **a**, Example one-shot learning task. The participant is asked to correctly identify a novel image (gray box) as an example of either object a (blue box) or object b (green box), given one example of each. **b**, The participant is asked to indicate previous familiarity with each of the visual concepts to be tested. We will use this information to ensure that we are evaluating *novel* concept learning. **c**, We collect the predicted 1-shot learning errors on a proposed set of unfamiliar objects, obtained by performing 1-shot learning experiments on visual concept manifolds in a trained ResNet50. The pattern of errors exhibits a rich structure, and includes a number of visual concept pairs whose errors are dramatically asymmetric.

**Supplementary Figure 8:**
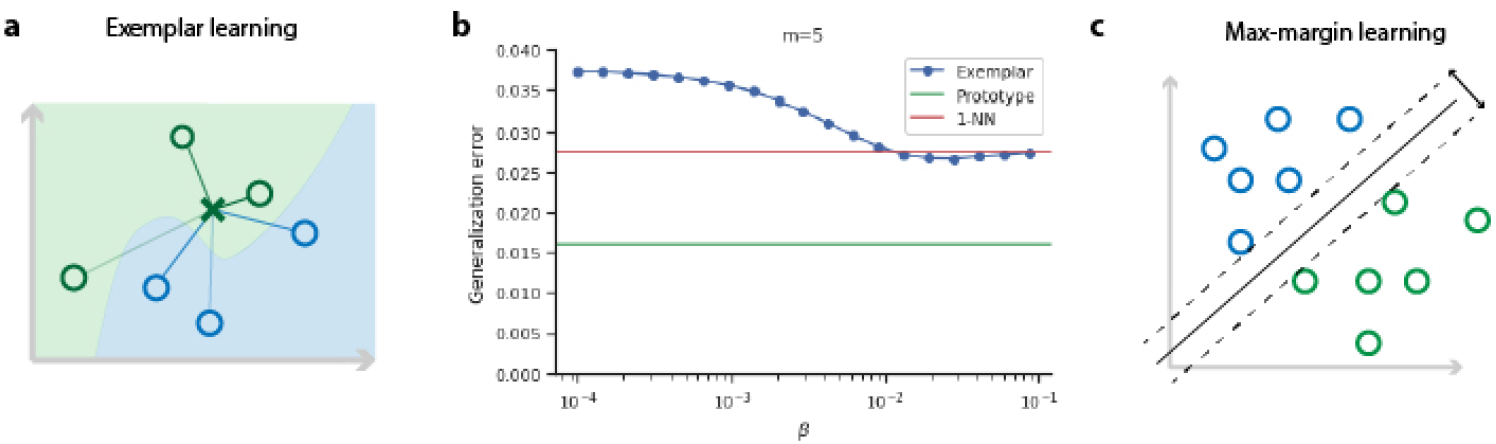
Comparing cognitive learning models. **a**, Under exemplar learning, a test example (green cross) is classified based on its similarity to each of the training examples (green and blue open circles). Hence exemplar learning involves the choice of a parameter *β* which weights the contribution of each training example to the discrimination function. When *β* = 0, all training examples contribute equally. When *β* = ∞, only the training example most similar to the test example contributes to the discrimination function. **b**, We perform exemplar learning experiments on concept manifolds in a trained ResNet50, and evaluate the generalization error as a function of *β*. We find that the optimal choice of *β* is large, approaching the *β* → ∞ limit. Furthermore, the optimal generalization error is very close to the *β* =∞ limit, which is equivalent to a nearest neighbors classifier (1-NN), whose generalization error is shown in red. For comparison, the generalization error of a prototype classifier is shown in green. **c**, Illustration of a max-margin classifier. The decision hyperplane (solid black line) of a max-margin classifier is optimized so that its minimum distance to each of the training examples is maximized^42^.

**Supplementary Figure 9:**
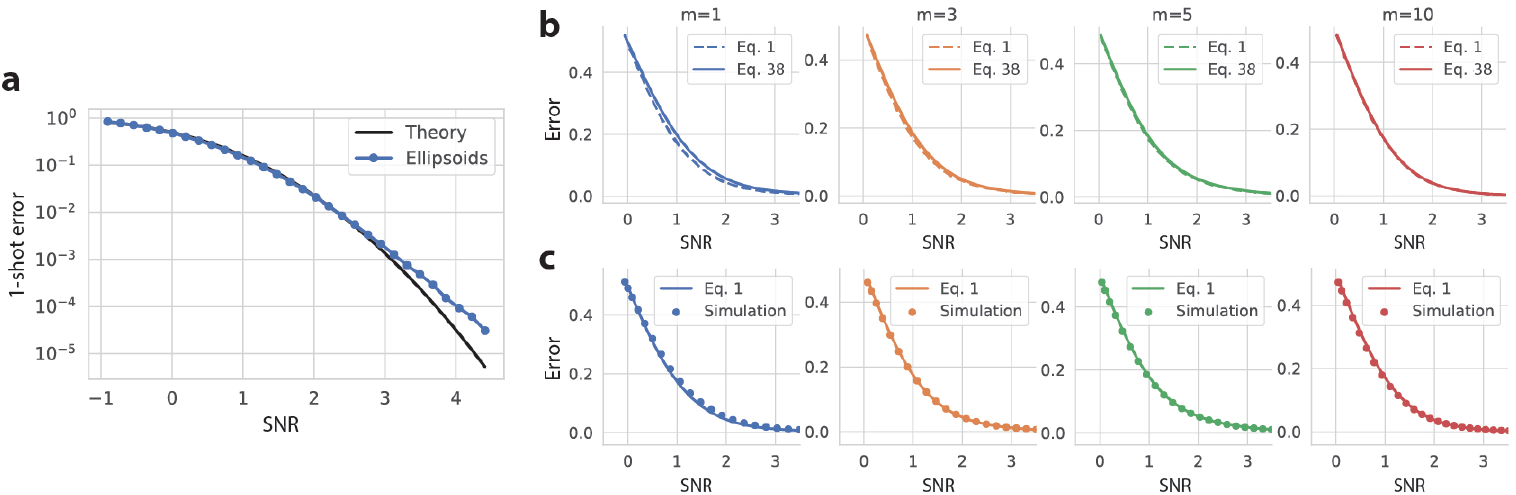
Numerical evaluation of the approximations used in our theory. **a**, Our theory for the few-shot learning SNR (see SI 2) approximates the projection of concept manifolds onto the linear readout direction as Gaussian-distributed. As discussed in SI 2.2, we expect this approximation to hold well when the SNR is small, and to break down when the SNR is large. To investigate the validity of this approximation, we perform numerical experiments on synthetic ellipsoids constructed to match the geometry of ImageNet21k visual concept manifolds in a trained ResNet50. For each pair of concept manifolds, we vary the signal ‖**Δ*x***_**0**_‖ ^2^ over the range 0.01 to 25 and perform 1-shot learning experiments. We compare the generalization error measured in experiments (blue points) to the prediction from our theory (Eq. SI.38; dark line). The theory closely matches experiment over several decades of error, and begins to break down for errors smaller than 10^−3^. Since errors smaller than 10^−3^ are difficult to resolve experimentally using real visual stimuli –as we have fewer than 1, 000 examples of each visual concept, and hence the generalization error may be dominated by one or a few outliers– we judge that this approximation holds well in the regime of interest. The match between theory and experiment for *m >* 1 shot learning (not shown) is as close or closer than for 1-shot learning, due to a law of large numbers-like effect. **b, c**, The few-shot learning SNR in the main text, Eq. 1, differs from the full SNR derived in SI 2.3, Eq. SI.38, which includes several additional terms. In **b** we investigate the difference between the two expressions. The two theoretical curves are nearly indistinguishable for *m* ≥ 3, but differ noticeably for *m* = 1. In **c** we compare Eq. 1 to the empirical generalization error measured in few-shot learning experiments on synthetic concept manifolds constructed to match the geometry of ImageNet21k visual concept manifolds in a trained ResNet50. The theory closely matches experiments for *m* ≥ 3, but slightly underestimates the generalization error for *m* = 1.

## Supplementary Materials

### 1 Introduction

In this supplementary material we develop our geometric theory for the generalization error of few-shot learning of high-dimensional concepts, we fill in the technical details associated with the main manuscript, and we perform more detailed investigations extending the results we have introduced. The outline of the supplementary material is as follows.

In SI 2 we derive an analytical prediction for the generalization error of prototype learning. We begin with a brief review of prototype learning using neural representations (SI 2.1). We then derive an exact expression for the generalization error of concept learning in a simplified model (SI 2.2), before proceeding to the full theory on pairs of novel concepts (SI 2.3). We then extend our model and theory to capture learning of more than two novel concepts in SI 2.4.

In SI 3 we examine the task of learning novel visual concepts without visual examples (zero-shot learning). We introduce a geometric theory for the generalization error of zero-shot learning in SI 3.1. We then compare the performance of zero-shot learning to few-shot learning, examining the question *how many words is an image worth?*, and identifying intriguing differences between the geometry of language-derived prototypes and vision-derived prototypes that govern the relative performance of the two models (SI 3.2).

In SI 4 we derive analytical predictions, drawing on the theory of random projections, for the number of neurons that must be recorded to reliably measure concept manifold geometry (SI 4.1), as well as the number of IT-like neurons a downstream neuron must listen to in order to achieve high few-shot learning performance (SI 4.2). In SI 5 we compare the performance of two foundational cognitive learning rules: prototype and exemplar learning, and we derive a fundamental relationship between concept dimensionality and the number of training examples that governs the relative performance of the two models.

In SI 6 we investigate the rich semantic structure encoded in the geometry of concept manifolds in trained DNNs. We show that the tree-like semantic organization of visual concepts in the ImageNet dataset is reflected in the geometry of visual concept manifolds, and that few-shot learning accuracy on pairs of novel concepts increases with the distance between the two concepts on the semantic tree, due to changes in each of the four geometric quantities identified in our theory. We additionally quantify the effect of distribution shift between the familiar concepts used to train the DNN, and novel concepts used to evaluate few-shot learning performance.

### 2 A geometric theory of few-shot learning

#### 2.1 Prototype learning using neural representations

Our model posits that novel concepts can be learned by learning to discriminate the manifolds of neural activity they elicit in higher order sensory areas, such as IT cortex. We further posit that learning can be accomplished by a population of downstream neurons via a simple plasticity rule. In the following sections we will introduce an analytical theory for the generalization error of concept learning using a particularly simple and biologically plausible plasticity rule: prototype learning. However, we find that this theory also correctly predicts the generalization error of more complex plasticity rules which involve learning a linear readout, such as max-margin learning, when concept manifolds are high-dimensional and the number of training examples is small. Furthermore, when concept manifolds are high-dimensional, their projection onto the linear readout direction is approximately Gaussian, and well characterized by the mean and covariance structure of the concept manifolds. For this reason we approximate concept manifolds as high-dimensional ellipsoids. We find that this approximation predicts the generalization error of few-shot learning remarkably well, despite the obviously complex shape of concept manifolds in the brain and in trained DNNs.

#### 2.2 Exact theory for high-dimensional spheres in orthogonal subspaces

Before proceeding to the full theory, we begin by studying a toy problem which simplifies the analysis and highlights some of the interesting behavior of few-shot learning in high dimensions. We examine the problem of classifying two novel concepts whose concept manifolds are high-dimensional spheres. Each sphere can be described by its centroid 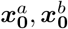, and its radius *R*_*a*_, *R*_*b*_, along a set of orthonormal axes 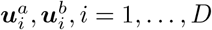, where we assume that each manifold occupies a *D*-dimensional subspace of the *N* -dimensional firing rate space. We will further assume that these subspaces are mutually orthogonal, 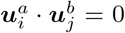, and orthogonal to the centroids, 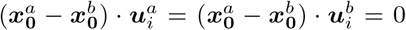, so that the signal-noise overlaps are zero. Thus a random example from each manifold can be written as,

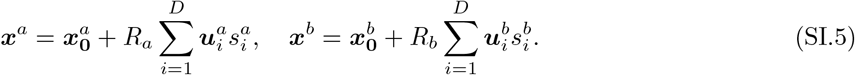

where ***s***^*a*^, ***s***^*b*^ ∼ Unif(𝕊^*D*−1^) are random vectors sampled uniformly from the *D*-dimensional unit sphere. We will study 1-shot learning in this section, using ***x***^*a*^, ***x***^*b*^ as training examples to learn a decision rule, and proceed to few-shot learning in the next section. Notice that in the 1-shot setting, prototype learning, max-margin learning, and exemplar learning all correspond to the same decision rule, which simply categorizes a test example of concept *a*, ***ξ***^*a*^, based on whether it is more similar to ***x***^*a*^ or ***x***^*b*^. Hence the theory we derive in this section is general to prototype learning, max-margin learning, and exemplar learning, as well as a wide range of other learning rules. The test example ***ξ***^*a*^ can be written as,

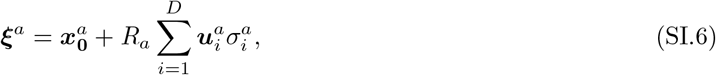

where ***σ***^*a*^ *∼* Unif(𝕊^*D*−1^) is a random vector sampled uniformly from the *D*-dimensional unit sphere. Using the Euclidean distance metric, ***ξ***^*a*^ is classified correctly if 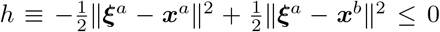. This decision rule corresponds to a linear classifier, and can be implemented by a downstream neuron which adjusts its synaptic weight vector ***w*** to point along the difference between the training examples, ***w*** = ***x***^*a*^ − ***x***^*b*^, and adjusts its firing threshold (bias) *β* to equal the average overlap of ***w*** with each training example, *β* = ***w*** · (***x***^*a*^ + ***x***^*b*^)*/*2. Then the output of the linear classifier on a test example ***ξ***^*a*^ is 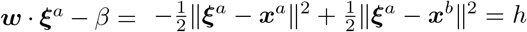, which can be thought of as the membrane potential of the downstream neuron. The generalization error on concept *a, ε*_*a*_, is given by the probability that this test example is incorrectly classified, *ε*_*a*_ = ℙ [*h*≤ 0]. Evaluating *h* using our parameterizations for ***x***^*a*^, ***x***^*b*^, ***ξ***^*a*^ (Eqs. SI.5, SI.6) gives,

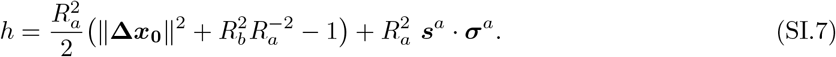

Where we have defined 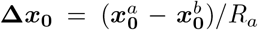 Thus we can evaluate the generalization error by computing *ε*_*a*_ = ℙ [*h ≤* 0] over all draws of the training and test examples. Defining 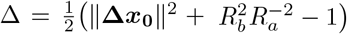,

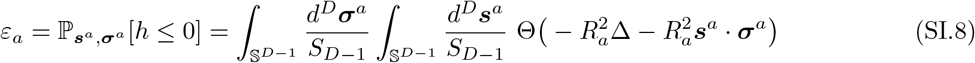

where Θ(·) is the Heaviside step function, and *S*_*D*−1_ is the surface area of the *D*-dimensional unit sphere. Enforcing the spherical constraint via a delta function,

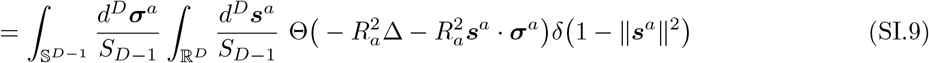

Writing the delta and step functions using their integral representations,

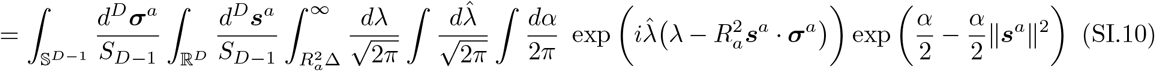

We now perform the Gaussian integral over *s*^*a*^,

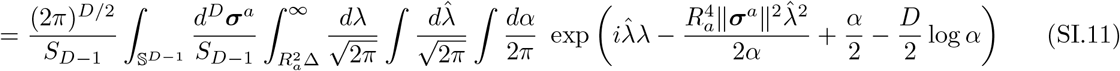

Noting that ‖***σ***^*a*^‖^2^ is constant over the unit sphere, the integral over ***σ***^*a*^ drops out,

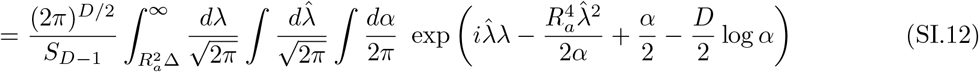

Performing the Gaussian integral over 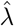,

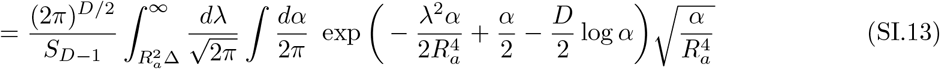

Expressing the result in terms of the Gaussian tail function 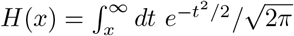,

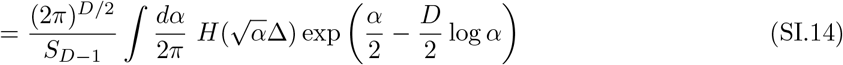

We evaluate the integral over *α* by saddle point. The saddle point condition is,

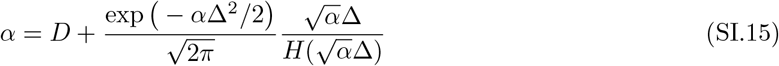

We will begin by studying the case where 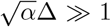, and revisit the case where 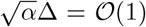. When 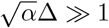, solving for *α* gives

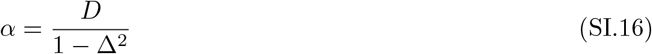

Noting that *S*_*D*−1_ is similarly given by *S*_*D*−1_ =∫*dα* ′ exp(*α* ′/2 − *D* log(*α* ′)(2*π*)^*D*/2^,we obtain the saddle point condition *α* ′= *D*. Using these conditions, we evaluate the integral in Eq. SI.14 at the saddle point, yielding,

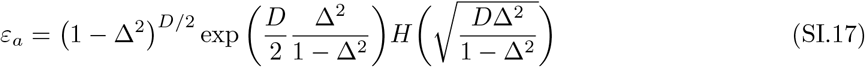

This expression reveals a sharp zero-error threshold at Δ = 1, reflecting a geometric constraint due to the bounded support of each spherical manifold. The generalization error is strictly zero whenever 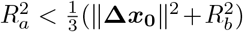. However, when *D* is large, the generalization error becomes exponentially small well before this threshold, when Δ *≪* 1 and 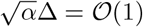. Indeed, the generalization error of prototype learning on concept manifolds in DNNs and macaque IT is better described by the regime where 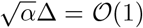. In this regime, the saddle point condition (Eq. SI.15) gives *α* = *D*, and the generalization error takes the form,

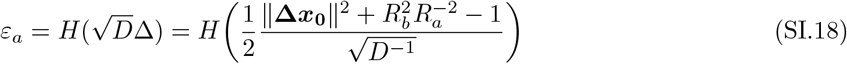

Hence in this regime the generalization error is governed by a signal-to-noise ratio which highlights some of the key behavior of the full few-shot learning SNR (Eq. 1). First, the SNR increases with the separation between the concept manifolds ‖**Δ*x***_**0**_ **‖**^2^. Second, the SNR increases as the manifold dimensionality *D* increases. As Fig. 2**c** shows, this is due to the fact that the projection of each manifold onto the linear readout direction ***w*** concentrates around its mean for large *D*. Remarkably, no matter how close the manifolds are to one another, the generalization error can be made arbitrarily small by making *D* sufficiently large. Third, the generalization error depends on an asymmetric term arising from the classifier bias, 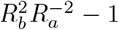. Decreasing *R*_*b*_ for fixed *R*_*a*_ increases *ε*_*a*_, while increasing *R*_*b*_ for fixed *R*_*a*_ decreases *ε*_*a*_. Interestingly, increasing *R*_*b*_ beyond 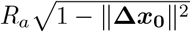 yields a *negative* SNR, and hence a generalization error worse than chance.

The dependence of Eq. SI.18 on the Gaussian tail function *H*(·) suggests that the projection of the concept manifold onto the readout direction ***w*** is well approximated by a Gaussian distribution. This approximation holds when the SNR is 𝒪 (1), but breaks down when the SNR is large. Motivated by the observation that the few-shot learning SNR for concept manifolds in macaque IT and DNNs is 𝒪 (1) (Figs. 4,5), we will use this approximation in the following section to obtain an analytical expression for the generalization error in the more complicated case of ellipsoids in overlapping subspaces, for which no exact closed form solution exists. We investigate the validity of this approximation quantitatively in Supp. Fig. 9**a**. We perform few-shot learning experiments on synthetic ellipsoids constructed to match the geometry of ResNet50 concept manifolds, and compare the empirical generalization error to the theoretical prediction derived under this approximation. Theory and experiment match closely for errors greater than 10^−3^. Since errors smaller than 10^−3^ are difficult to resolve experimentally using real visual stimuli –as we have fewer than 1, 000 examples of each visual concept, and hence the generalization error may be dominated by one or a few outliers– we judge that this approximation holds well in the regime of interest.

#### 2.3 Full theory: high-dimensional ellipsoids in overlapping subspaces

We now proceed to the full theory for few-shot learning on pairs of high-dimensional ellipsoids, relaxing the simplifying assumptions in the previous section. We draw *µ* = 1, …, *m* training examples each from two concept manifolds, *a* and *b*,

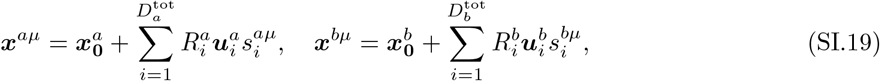

Where 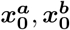 are the manifold centroids, and 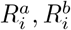 are the radii along each axis, 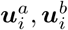. 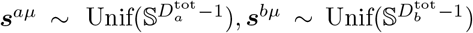are random samples from the unit sphere. 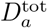 and 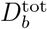 represent the total number of dimensions along which each manifold varies. In practical situations 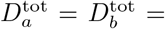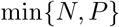, where *N* is the number of recorded neurons and *P* is the number of examples of each concept. To perform prototype learning, we average these training examples into prototypes, 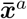 and 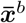,

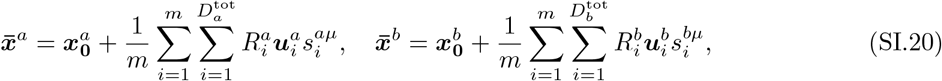

To evaluate the generalization error of prototype learning, we draw a test example

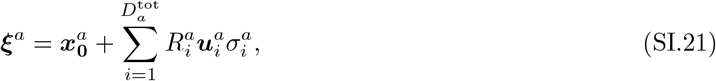

and compute the probability that ***ξ***^*a*^ is correctly classified, 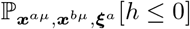, where 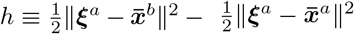 .Evaluating *h* using our parameterization gives,

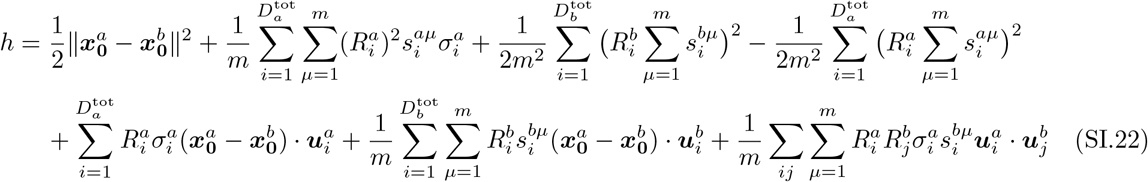

As we will see, the first term corresponds to the signal, the second to the dimension, the third and fourth terms to the bias, the fifth and sixth to signal-noise overlaps, and the seventh to noise-noise overlaps, which quantify the overlap between manifold subspaces. Each of these terms is independent and, as discussed in the previous section, approximately Gaussian-distributed when the dimensionality of concept manifolds is high. Hence by computing the mean and variance of each term we can estimate the full distribution over *h*. Noting that 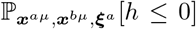 is invariant to an overall scaling of *h*, we will define the renormalized 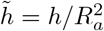, which is dimensionless. Computing the generalization error in terms of 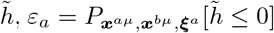, will allow us to obtain an expression which depends only on interpretable, dimensionless quantities.

##### Signal

The first term in Eq. SI.22, corresponding to signal, is fixed across different draws of the training and test examples, and so has zero variance. Its mean is given by 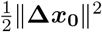, where 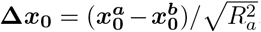.

##### Dimension

The second term in Eq. SI.22 corresponds to the manifold dimension. Its mean is zero, since by symmetry odd powers of 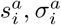 integrate to zero over the sphere. Quadratic terms integrate to 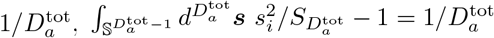; hence the variance is given by,

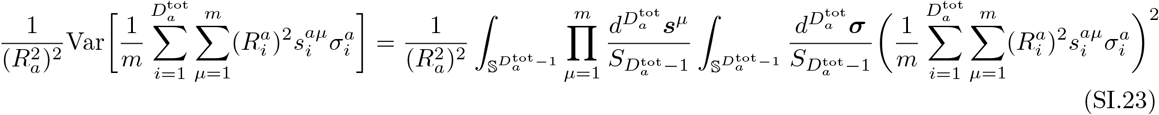

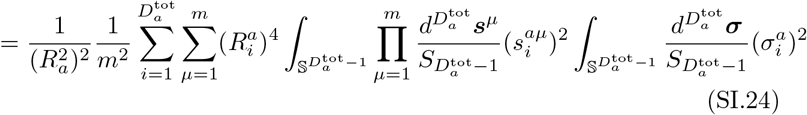

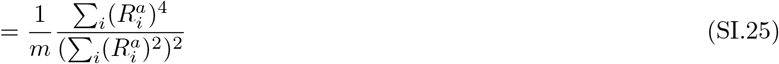

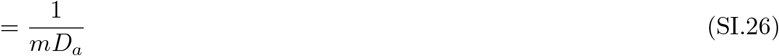

Where 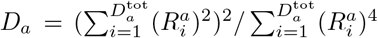is the participation ratio, which measures the effective dimensionality of the concept manifold, quantified by the number of dimensions along which it varies significantly^29^. Hence this term reflects the manifold dimensionality, and its variance is suppressed for large *D*_*a*_.

##### Bias

We next proceed to the third and fourth terms of Eq. SI.22, which correspond to bias. We show only the calculation for the first bias term, as the second bias term follows from the same calculation. The mean is given by,

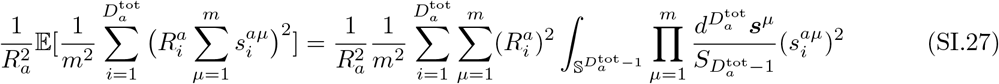

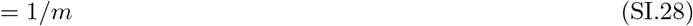

And the variance is given by,

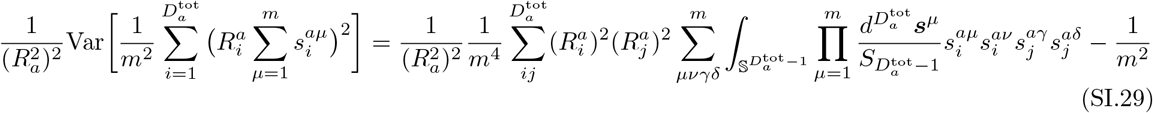

There are three possible pairings of indices which yield even powers of *s*_*i*_. Due to symmetry, all other pairings integrate to zero. First, there are *m* terms of the form 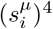, each of which integrates to 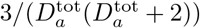. Second, there are 3*m*(*m* −1) terms of the form 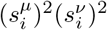, each of which integrates to 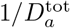. Finally, there are *m*^2^ terms of the form 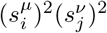, each of which integrates to 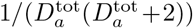 .Thus the integral gives,

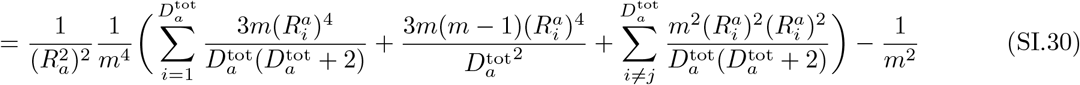

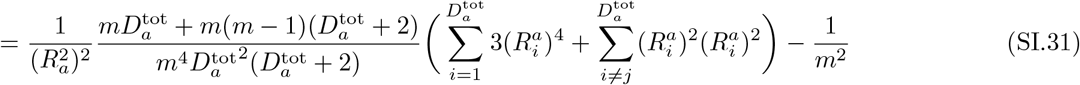

Dropping small terms of 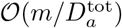, and writing the final expression in terms of the effective dimensionality *D*_*a*_,

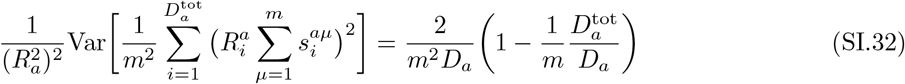

Notice that when *m* = 1 and the radii are spread equally over all dimensions, so that 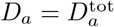 (i.e. the manifold is a sphere), the variance goes to zero. However, in practical situations the effective dimensionality is much smaller than the total number of dimensions, 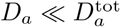, and the variance is given by 2*/m*^2^*D*_*a*_.

##### Signal-noise overlaps

We now proceed to the signal-noise overlap terms on the second line of Eq. SI.22, each of which has zero mean. The variance of the first signal-noise overlap term is given by,

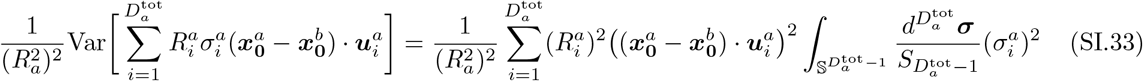

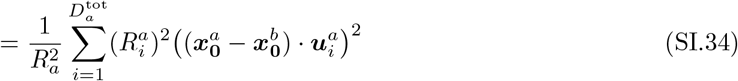

We refer to this term as signal-noise overlap because it quantifies the overlap between the noise directions 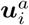, and the signal direction **Δ*x***_**0**_, weighted by the radii 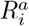 along each noise direction. To make the notation more compact, we define 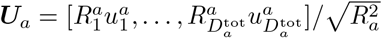, so that the signal-noise overlap takes the form,

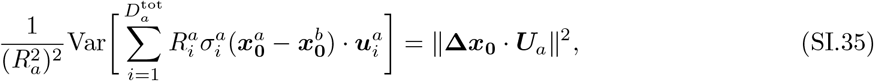

Notice that this signal-noise overlap term does not depend on *m*, since it involves only the test examples. The second signal-overlap term, in contrast, captures the variation of the training examples along the signal direction, and so its variance does depend on *m*,

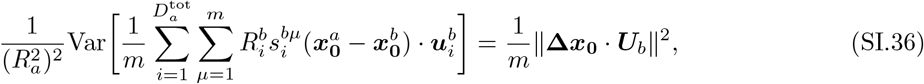

where we have defined 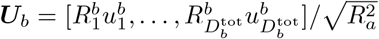 in analogy to ***U***_*a*_. As the number of training examples increases, the variation of the *b* prototype along the signal direction decreases, and the contribution of this signal-noise overlap term decays to zero.

##### Noise-noise overlaps

Finally, we compute the mean and variance of the final term of Eq. SI.22, the noise-noise overlap term, which follows from a similar calculation. The mean is given by zero, and the variance by,

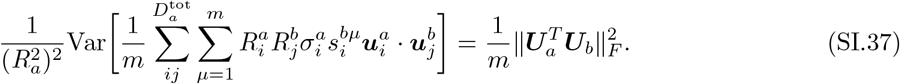

We refer to this term as the noise-noise overlap because it quantifies the overlap between the noise directions of manifold *a*, ***U***_*a*_, and the noise directions of manifold *b*, ***U***_*b*_.

##### SNR

Combining the terms computed above, the mean and variance of 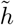 are given by,

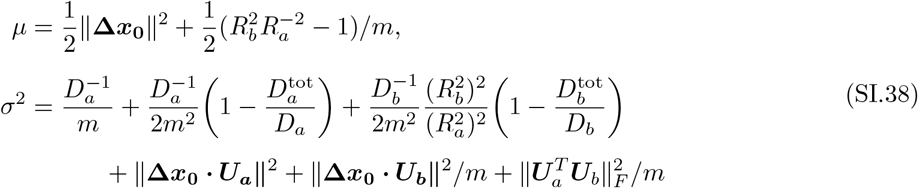

We will refer to the mean as the signal, and the standard deviation as the noise. Hence the generalization error can be expressed in terms of the ratio of the signal to the noise, 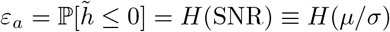. Suppressing terms in Eq. SI.38 which we argue contribute only a small correction yields the few-shot learning SNR in the main text, Eq. 1. These additional terms, whose contribution we quantify in Supp. Fig. 9**b**,**c**, are the two noise terms arising from the bias,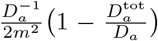 and 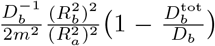,and the noise-noise overlaps term 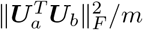. We find that for concept manifolds in macaque IT and in DNN concept manifolds, noise-noise overlaps are substantially smaller than signal-noise overlaps and 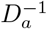, and their contribution to the overall SNR is negligible. The two noise terms arising from the bias fall off quadratically with *m*, and we find that their contribution is negligible for *m*≥ 3 (Supp. Fig. 9**b**,**c**). Indeed, by performing few-shot learning experiments using synthetic ellipsoids constructed to match the geometry of ImageNet21k visual concept manifolds in a trained ResNet50 (Supp. Fig. 9**b**), we find that Eq. 1 and Eq. SI.38 are nearly indistinguishable for *m* ≥ 3. However, for *m* = 1 the additional terms in Eq. SI.38 yield a small but noticeable correction. Consistent with this, we find that Eq. 1 accurately predicts the empirical generalization error measured in few-shot learning experiments for *m*≥ 3, but very slightly underestimates the generalization error for *m* = 1 (Supp. Fig. 9**c**). For this reason we include only the dominant terms in the main text (Eq. 1), but we use Eq. SI.38 to predict the generalization error in simulations when *m ≤* 3.

#### 2.4 Learning many novel concepts from few examples

Concept learning often involves categorizing more than two novel concepts (Supp. Fig. 2**a**). Here we extend our model and theory to the case of learning *k* new concepts, also known as *k*-way classification. Prototype learning extends naturally to *k*-way classification: we simply define *k* prototypes, 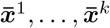, by averaging the training examples of each novel concept (Supp. Fig. 2**b**). A test example ***ξ***^*a*^ of concept *a* is classified correctly if it is closest in Euclidean distance to the prototype 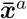 of concept *a*. That is, if *h*_*b*_ *>* 0 for all *b* ≠ *a*, where

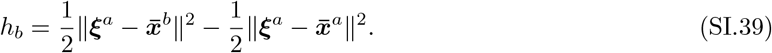

Notice that *h*_*b*_ can be rewritten as 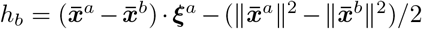. Hence this classification rule is linear, and can be implemented by *k* downstream neurons, one for each novel concept. Each downstream neuron adjusts its synaptic weight vector ***w***^*b*^ to point along the direction of a concept prototype, 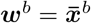, *b* = 1, …, *k*, and adjusts its firing threshold (bias) *β* to equal the overlap of ***w***^*b*^ with the prototype, 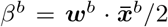. Then the test example ***ξ***^*a*^ of concept *a* is classified correctly if the output of neuron *a*, ***w***^*a*^ · ***ξ***^*a*^ − *β*^*a*^, is greater than the output of neuron *b*, ***w***^*b*^ · ***ξ***^*a*^ − *β*^*b*^, for all *b* ≠ *a*.

The generalization error on concept *a, ε*_*a*_, is given by the probability that at least one *h*_*b*_ *≥* 0, for all *b* ≠ *a*. Equivalently,

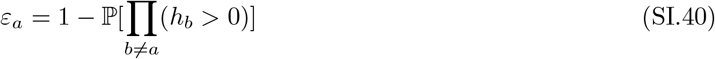

To evaluate this probability, we consider the joint distribution of the *h*_*b*_ for *b* ≠ *a*, defining the random variable ***h*** ≡ [*h*_1_, …, *h*_*a*−1_, *h*_*a*+1_, …, *h*_*k*_]. We have already computed *h*_*b*_ (Eq. SI.22) and seen that it is a Gaussian distributed random variable when the SNR= 𝒪 (1) and the concept manifold is high-dimensional. Hence in this regime ***h*** is distributed as a multivariate Gaussian random variable,

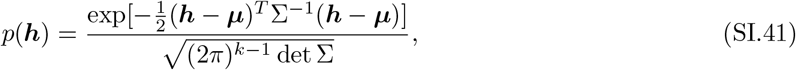

with mean *µ*_*b*_ *≡*𝔼 [*h*_*b*_], and covariance Σ_*bc*_ = 𝔼 [*h*_*b*_*h*_*c*_] − *µ*_*b*_*µ*_*c*_. We can therefore obtain the generalization error by integrating *p*(***h***) over the positive orthant, where all *h*_*b*_ *≥* 0,

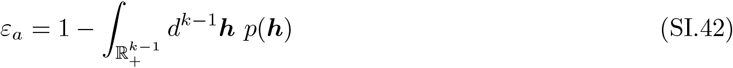

All that is left to do is compute the mean ***µ*** and covariance Σ. As before, ℙ [Π_*b≠a*_(*h*_*b*_ *>* 0)] is invariant to an overall scaling of *h*_*b*_, so we will work with the renormalized 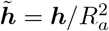 in order to obtain dimensionless quantities. We have already evaluated the mean 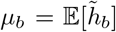 and the diagonal covariance elements 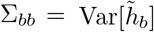 in SI 2.3; these are just the signal and noise, respectively, from the two-way SNR, Eq. SI.38. So we proceed to the off-diagonal covariances, 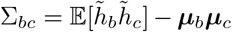. Using the expression for *h*_*b*_ in Eq. SI.22, we find that when *b* ≠ *c* three terms contribute,

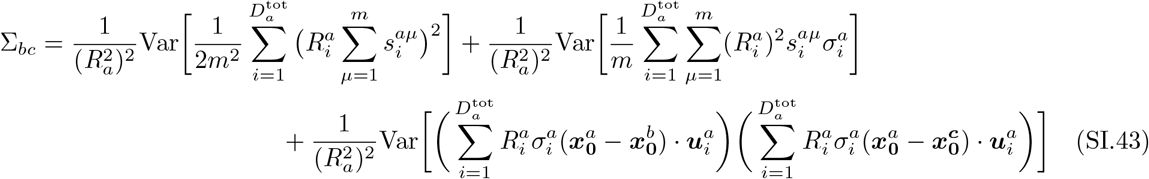

The first term we evaluate in Eq. SI.32,

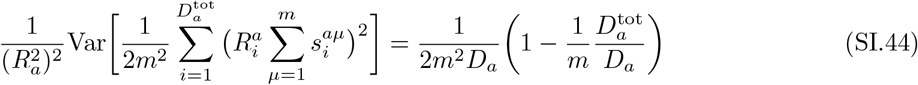

The second term we evaluate in Eq. SI.26,

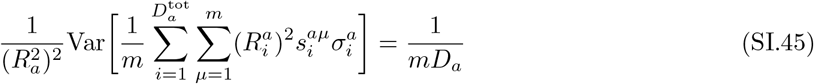

And for the third term we evaluate an analogous expression in Eq. SI.35, yielding,

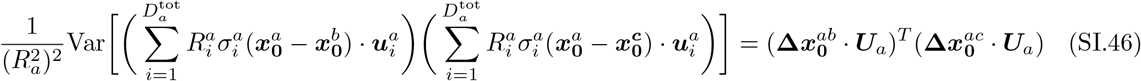

where 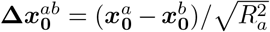, and 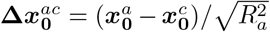. Combining these terms, and re-inserting the terms for *b* = *c* derived in Eq. SI.38, we obtain the full expression for the covariance,

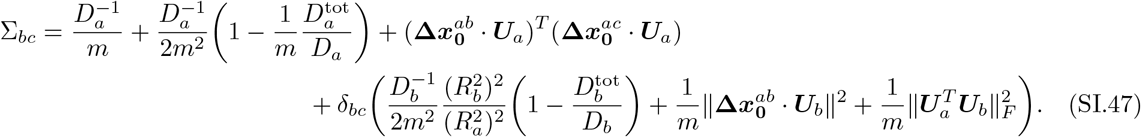

Recall from Eq. SI.38 that ***µ*** is given by,

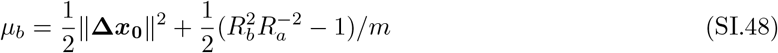

Integrating the multivariate Gaussian with mean ***µ*** and covariance Σ over the positive orthant, Eq. SI.42, gives the generalization error (Supp. Fig. 2).

### 3 Learning visual concepts without visual examples by aligning language to vision

#### 3.1 A geometric theory of zero-shot learning

Prototype learning also extends naturally to the task of learning novel visual concepts without visual examples (*zero-shot* learning), as we demonstrate in Section 2.7 by generating visual prototypes from language-derived representations. Moreover, our theory extends straightforwardly to predict the performance of zero-shot learning in terms of the geometry of concept manifolds. Consider the task of learning to classify two novel visual concepts, given concept prototypes ***y***^*a*^, ***y***^*b*^ derived from language, or from another sensory modality. To classify a test example of concept *a*, we present the test example to the visual pathway and collect the pattern of activity ***ξ***^*a*^ it elicits in a population of IT-like neurons. We then classify ***ξ***^*a*^ according to which prototype it is closer to. As in few-shot learning, we assume that ***ξ***^*a*^ lies along an underlying ellipsoidal manifold,

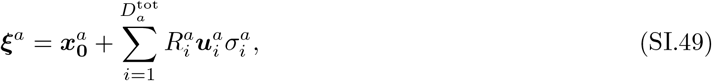

where 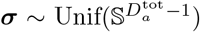. We define 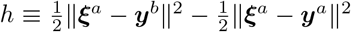, so that the generalization error is given by the probability that *h ≤* 0, 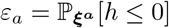. Writing out *h*,

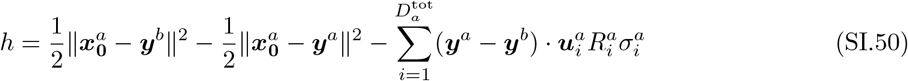

Hence the error depends only on the distances between the prototypes and the true manifold centroids, and the overlap between the manifold subspace and the difference between the two prototypes. When the concept manifold is high dimensional, the last term is approximately Gaussian-distributed, with zero mean and variance,

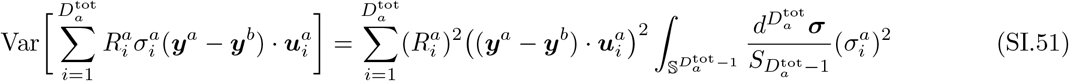

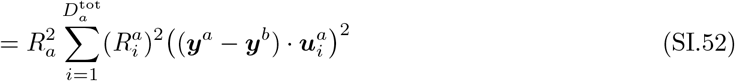

Defining 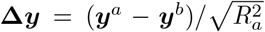 the variance can be written more compactly as 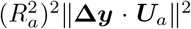.Hence the generalization error of zero-shot learning is governed by a signal-to-noise ratio, 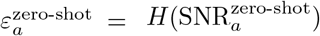,

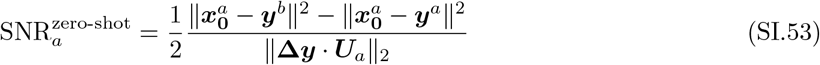

Where we have normalized all quantities by 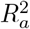. This theory yields a close match to zero-shot learning experiments performed on concept manifolds in a trained ResNet50 (Fig. 7**d**), and affords deeper insight into the performance of zero-shot learning, as we show in Fig. 7**e**, and explore further in the following section.

#### 3.2 many words is a picture worth? Comparing prototypes derived from language and vision

We found that prototypes derived from language yield a better generalization accuracy than those derived from a single visual example (Section 2.7), but not two or more visual examples (Supp. Fig. 4**a**). To better understand this behavior, we use our geometric theory for zero-shot learning, Eq. 3, to decompose the zero-shot learning SNR into a contribution from the ‘signal’, which quantifies how closely the estimated prototypes match the true manifold centroids, and a contribution from the ‘noise’, which quantifies the overlap between the readout direction and the noise directions. We use the same theory to examine the prototypes generated by few-shot learning, even though these prototypes vary across different draws of the training examples, by averaging the signal and noise over many different draws of the training examples. This allows us to compare zero-shot learning and few-shot learning in the same framework, to understand whether the enhanced performance of zero-shot learning is due to higher signal (i.e. a closer match between estimated prototypes and true centroids) or lower noise (i.e. less overlap between the readout and noise directions). In Supp. Fig. 4**b**,**c** we show that both signal and noise are significantly lower for zero-shot learning than for few-shot learning. Therefore, one-shot learning prototypes more closely match the true concept prototypes on average than language prototypes do. However, language prototypes are able to achieve a higher overall generalization accuracy by picking out linear readout directions which overlap significantly less with the concept manifolds’ noise directions. We visualize these directions in Supp. Fig. 4**d** by projecting pairs of concept manifolds into the two-dimensional space spanned by the signal direction **Δ*x***_**0**_ and the language prototype readout direction **Δ*y***. In each case, the manifolds’ variability is predominantly along the signal direction **Δ*x***_**0**_, while the language prototypes pick out readout directions **Δ*y*** with much lower variability.

### 4 How many neurons are required for concept learning?

Neurons downstream of IT cortex receive inputs from only a small fraction of the total number of available neurons in IT. How does concept learning performance depend on the number of input neurons? Similarly, a neuroscientist seeking to estimate concept manifold geometry in IT only has access to a few hundred neurons. How is concept manifold geometry distorted when only a small fraction of neurons is recorded from?

In this section we will draw on the theory of random projections to derive analytical answers to both questions. We will model recording from a small number *M* of the *N* available neurons as projecting the *N* -dimensional activity patterns into an *M* -dimensional subspace. When activity patterns are randomly oriented with respect to single neuron axes, selecting a random subset of neurons to record from is exactly equivalent to randomly projecting the full *N* -dimensional activity patterns into an *M* -dimensional subspace. We will begin by deriving the behavior of concept manifold dimensionality *D* as a function of the dimension of the target space *M*, and use this to derive the behavior of the few-shot learning generalization error.

#### 4.1 Concept manifold dimensionality under random projections

Consider randomly projecting each point ***x*** ∈ ℝ^*N*^ on a concept manifold to a lower-dimensional subspace, *A****x*** = ***x′*** ∈ ℝ^*M*^ using a random projection matrix *A* ∈ ℝ^*M ×N*^, *A*_*ij*_ *∼𝒩* (0, 1*/M*). We collect all points on the original concept manifold into an *N × P* matrix *X*, and collect all points on the projected concept manifold into an *M × P* matrix *X* ′ = *AX*. Recall that the effective dimensionality *D*(*N*) of the original concept manifold can be expressed in terms of its *N × N* covariance matrix 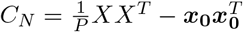,

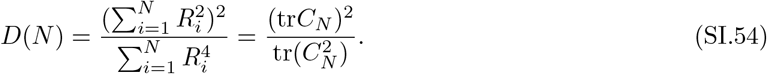

Likewise, the effective dimensionality *D*(*M*) of the projected concept manifold can be expressed in terms of its *M × M* covariance matrix 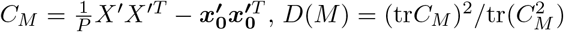.Notice that

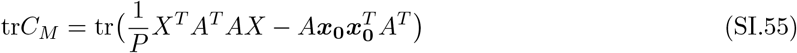

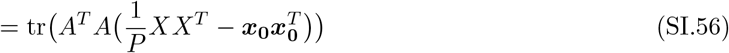

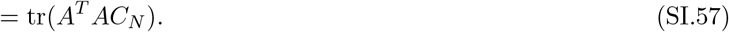

Where we have used the cyclic property of the trace. Hence the relationship between tr*C*_*N*_ and tr*C*_*M*_ is governed by the statistics of Λ ≡ *A*^*T*^ *A*. Λ is a Wishart random matrix, with mean 𝔼 [Λ] = *I* and variance Var[Λ] = 1*/M* + *I/M*. To estimate the effective dimensionality *D*(*M*) of the projected concept manifold, we can compute the expected value of (tr*C*_*M*_)^2^ and tr(*C*_*M*_)^2^ over random realizations of Λ.

We will start with the denominator of *D*(*M*), tr(*C*_*M*_)^2^,

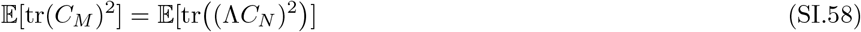

Diagonalizing *C*_*N*_ = *UR*^2^*U*^*T*^,

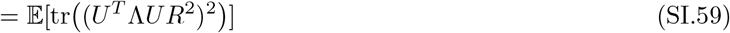

Defining 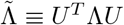,

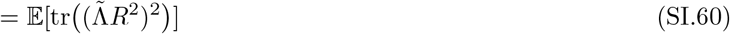

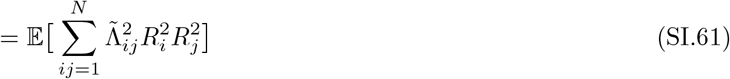

Notice that 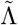 has the same statistics as Λ. Hence,

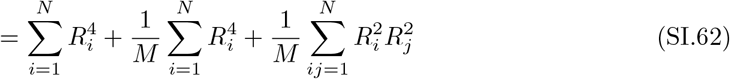

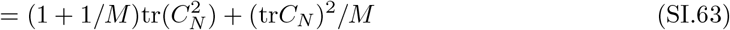

We now proceed to the numerator (tr*C*_*M*_)^2^,

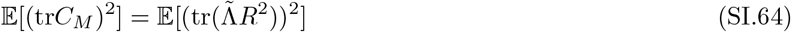

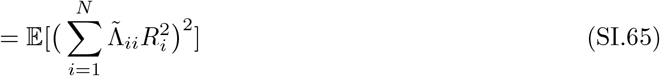

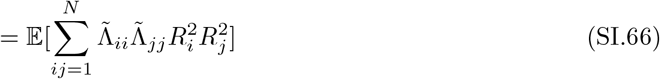

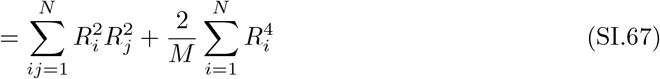

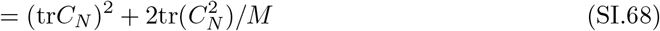

Combining our expressions for the numerator and the denominator, we obtain an estimate for the expected value of *D*(*M*),

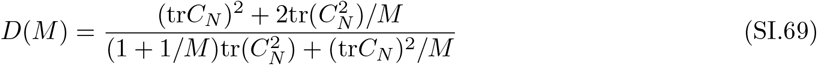

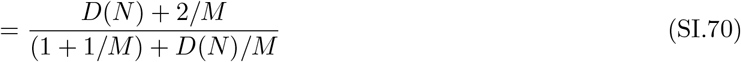

Dropping the small terms of order 1*/M*,

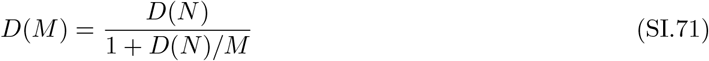

Therefore, provided that *M* is large compared to *D*, the random projection will have a negligible effect on the dimensionality. However, when *M* is on the order of *D*, distortions induced by the random projection will increase correlations between points on the manifold, significantly decreasing the effective dimensionality. Taking *N → ∞*, this expression also allows us to extrapolate the asymptotic dimensionality *D*_*∞*_ = *D*(*M*)*/*(1 −*D*(*M*)*/M*) we might observe given access to arbitrarily many neurons. When concept manifolds occupy only a small fraction of the *M* available dimensions given recordings of *M* neurons, then recording from a few more neurons will have only a marginal effect. But when concept manifolds occupy a large fraction of the *M* available dimensions, recording from a few more neurons may significantly increase the estimated manifold dimensionality. Using this asymptotic dimensionality *D*_*∞*_, we can obtain a single expression for the estimated dimensionality *D*(*M*) of concept manifolds given recordings of *M* neurons,

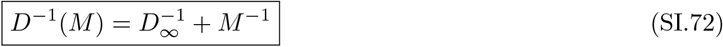

This prediction agrees well with random projections and random subsampling experiments on concept manifolds in IT and in trained DNNs (Fig. 6).

#### 4.2 Few-shot learning requires a number of neurons *M* greater than the concept manifold dimensionality *D*

We next ask how the generalization error of few-shot learning depends on the number of subsampled neurons. We will study the simple case of 1-shot learning on identical ellipsoids in orthogonal subspaces, and demonstrate empirically that the predictions we derive hold well for the full case. Recall that the 1-shot learning SNR for identical ellipsoids in orthogonal subspaces (SI 2.3) is given by

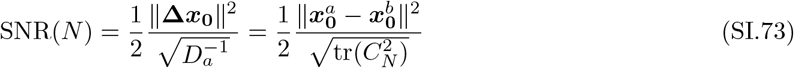

Then the signal-to-noise ratio in the projected subspace, SNR(*M*), is given by

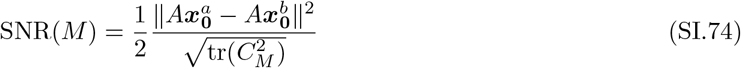

We have already found that 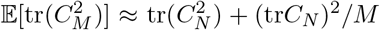. Furthermore, random projections are known to preserve the pairwise distances between high-dimensional points under fairly general settings, so that the distance between manifold centroids, 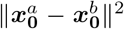, is preserved under the random projection, 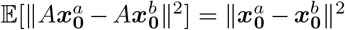. Deviations from this average are quantified by the Johnson-Lindenstrauss Lemma, a fundamental result in the theory of random projections, which states that *P* points can be embedded in *M* = *𝒪* (log *P/ϵ*^2^) dimensions without distorting the distance between any pair of points by more than a factor of (1 ± *ϵ*). Combining these results, we have

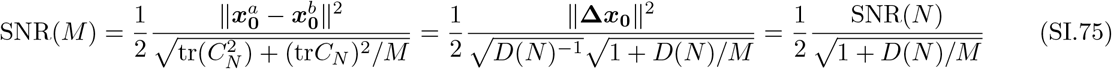

Therefore, few-shot learning performance is unaffected by the random projection, provided that *M* is large compared to the concept manifold dimensionality. As before, we can extrapolate an asymptotic SNR given access to arbitrarily many neurons by taking *N*→ ∞, 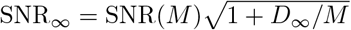. When concept manifolds occupy only a small fraction of the *M* available dimensions, a downstream neuron improves its few-shot learning performance only marginally by receiving inputs from a greater number of neurons. However, when concept manifolds occupy a large fraction of the *M* available dimensions, a downstream neuron can substantially improve its few-shot learning performance by receiving inputs from a greater number of neurons. Using this asymptotic signal-to-noise ratio, SNR_*∞*_, we can obtain a single expression for the few-shot learning SNR as a function of the number of input neurons, *M*,

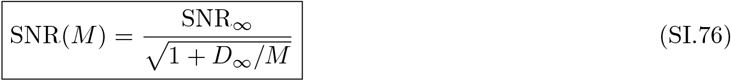

This prediction agrees well with random projections and random subsampling experiments on concept manifolds in IT and in trained DNNs (Fig. 6).

### 5 Comparing cognitive learning models in low and high dimensions

A long line of work in the psychology literature has examined the relative advantages and disadvantages of prototype and exemplar theories of learning. Exemplar learning is performed by storing the representations of all training examples in memory, and categorizing a test example by comparing it to each stored example (Supp. Fig. 8**a**). Exemplar learning thus involves a choice of how to weight the similarity to each of the training examples. In one extreme, all similarities are weighted equally, so that a test example is categorized as concept *a* if its average similarity to each of the training examples of concept *a* is greater than its average similarity to each of the training examples of concept *b*. This limit is analytically tractable, and we find that it performs consistently worse than prototype learning. Indeed, in our experiments the optimal weighting is very close to the opposite extreme, in which only the most similar training example is counted, and the test example is assigned to whichever category this most similar training example belongs to (Supp. Fig. 8**b**). This limit corresponds to a nearest-neighbor (NN) decision rule. In numerical experiments on visual concept manifolds in trained DNNs (Fig. 8**a**), we find that prototype learning outperforms NN when *D* is large and the number of training examples *m* is small, while NN outperforms prototype learning in the opposite regime where *D* is small and the number of training examples *m* is large. Here we offer a theoretical justification for this behavior. We begin with an intuitive summary, and proceed to a more detailed derivation in the following section.

#### 5.1 Identifying the joint role of dimensionality *D* and number of training examples *m*

The joint role of *D* and *m* arises because NN learning involves taking a minimum over the distances from each training example to the test example. However, as we have seen, in high dimensions these distances concentrate around their means with variance 1*/D*. Under fairly general conditions, the minimum over *m* independent random variables with variance 1*/D* scales as 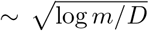. When all other geometric quantities are held constant, the signal of NN learning scales as 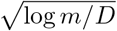, while the signal of prototype learning is constant. Hence when log *m* is large compared to *D*, NN learning outperforms prototype learning, and when *D* is large compared to log *m*, prototype learning outperforms NN learning.

We now derive the few-shot learning signal for NN learning, analogous to the few-shot learning signal we derived for prototoype learning, Eq. 1. The setup for NN learning is the same as for prototype learning: we draw *m* training examples each from two concept manifolds, *a* and *b*,

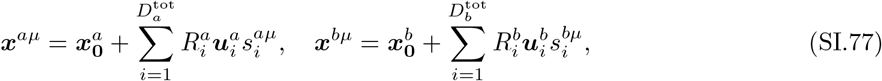

Where 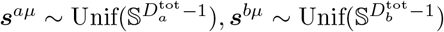. We then draw a test example,

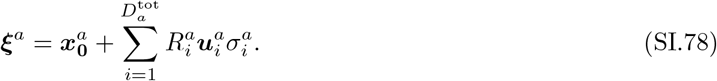

Where 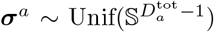. Rather than averaging the training examples into concept prototypes, to perform NN learning we simply compute the Euclidean distance from the test example to each of the training examples of concept *a*,

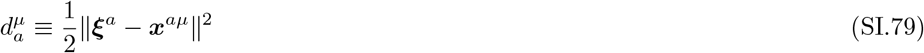

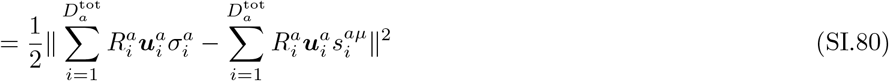

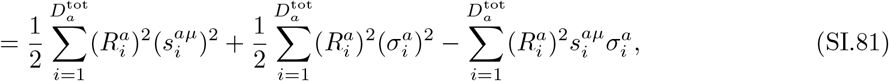

And the distance to each of the training examples of concept *b*,

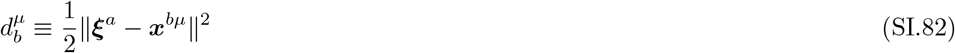

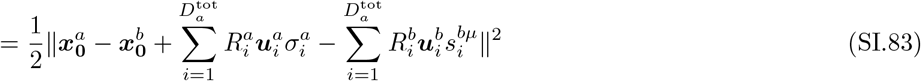

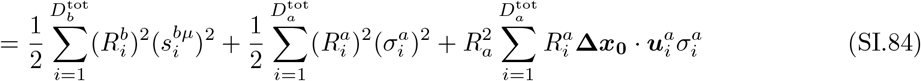

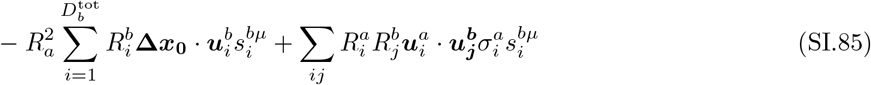

Then the generalization error is the probability that 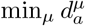 is less than 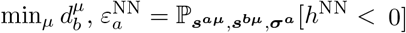, where 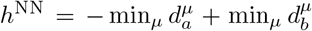. As we found in prototype learning, when concept manifolds are high-dimensional, 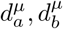 are approximately Gaussian-distributed. Again, in order to obtain dimensionless quantities we renormalize,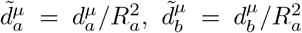.We define the mean 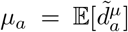 and variance 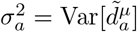, given by

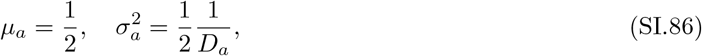

which follow from eqs. SI.27, SI.32, and SI.26. Similarly, we define 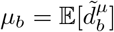 and 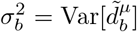, given by

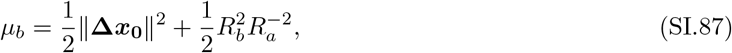

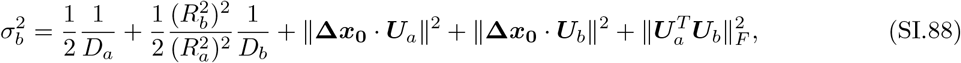

which follow from eqs. SI.27, SI.32, SI.35, and SI.37. Now we must evaluate the minimum over *µ*. The expected value of the minimum of *m* i.i.d. Gaussian random variables is given by 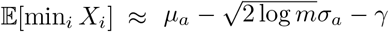 where 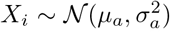, *i* = 1, …, *m* and *γ* is the Euler-Mascheroni constant. Using this we can obtain the expected value of 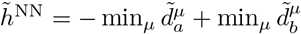,

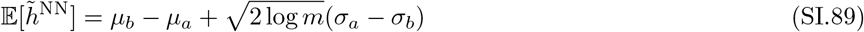

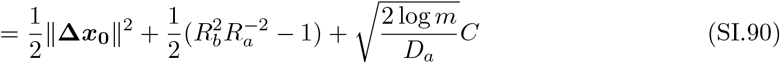

Where we have pulled the dependence on 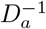 out of *σ*_*a*_, *σ*_*b*_ to define 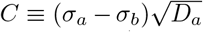 is greater than zero, since the signal-noise and noise-noise overlaps are much smaller than 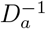, and therefore *σ*_*a*_ *> σ*_*b*_. Neglecting the bias term 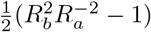, we have that the signal of prototype learning is given by

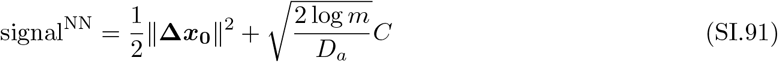

Compare this to the signal we found for prototype learning,

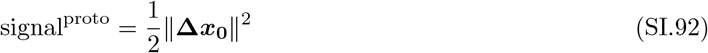

The NN signal is larger than the prototype learning signal. However the NN noise is also larger than the prototype learning noise. Hence when log *m* is large compared to *D*_*a*_, NN outperforms prototype learning, and when *D*_*a*_ is large compared to log *m*, protoype learning outperforms NN. We stop short of computing a full SNR for NN, since the random variables 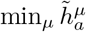 and 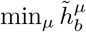 are not independent, and computing their correlation is not straightforward. However, the *D*∼ log *m* relationship we have identified here seems to reliably capture the behavior we observe in experiments on concept manifolds in a trained DNN (Fig. 8**a**), where we vary the dimensionality by projecting each concept manifold onto its top *D* principal components.

### 6 Geometry of DNN concept manifolds encodes a rich semantic structure

The ImageNet21k dataset is organized into a semantic tree, with each of the 1k visual concepts in our evaluation set representing a leaf on this tree (see Methods 3.1). To investigate the effect of semantic structure on concept learning, we sort the generalization error pattern of prototype learning in a trained ResNet50 to obey the structure of the semantic tree, so that semantically related concepts are adjacent, and semantically unrelated concepts are distant. The sorted error matrix (Supp. Fig. 1**a**) exhibits a prominent block diagonal structure, suggesting that most of the errors occur between concepts on the same branch of the semantic tree, and errors between two different branches of the semantic tree are exceedingly unlikely. In other words, the trained ResNet may confuse two types of Passerine birds, like songbirds and sparrows, but will almost never confuse a sparrow for a mammal or a fish. The sorted error matrix exhibits structure across many scales: some branches reveal very fine-grained discriminations (e.g. aquatic birds), while other branches reveal only coarser discriminations (e.g. Passerines). We suspect that the resolution with which the trained DNN represents different branches of the tree depends on the composition of the visual concepts seen during training, which we discuss further below. Finally, the sorted error pattern exhibits a pronounced asymmetry, with much larger errors above the diagonal than below. In particular, food and artifacts are more likely to be classified as plants and animals than plants and animals are to be classified as food and artifacts.

We additionally sort the patterns of individual geometric quantities: signal, bias, and signal-noise overlap, to reflect the semantic structure of the dataset (Supp. Fig. 1**a**, right). Signal exhibits a clear block diagonal structure, similar to the error pattern. Bias reveals a clear asymmetry: plants and animals have significantly higher bias than food and artifacts do, indicating that the radii of plant and animal concept manifolds are significantly smaller than the radii of food and artifact concept manifolds. Intriguingly, this suggests that the trained ResNet50 has learned more compact representations for plants and animals than for food and artifacts.

To quantify the extent to which each of these quantities depends on the semantic organization of visual concepts, we compute the average few-shot accuracy, signal, bias, and signal noise overlap across all pairs of concepts, as a function of the distance between the two concepts on the semantic tree, defined by the number of hops required to travel from one concept to the other (Supp. Fig. 1**b**). We find that few-shot learning accuracy, signal, and bias all increase significantly with semantic distance, while signal-noise overlaps decrease.

A related question is the effect of distribution shift between trained and novel concepts. The composition of the 1, 000 heldout visual concepts in our evaluation set is quite different from that of the 1, 000 concepts seen during training. For instance, 10% of the training concepts are different breeds of dogs, while only 0.5% of the novel concepts are breeds of dogs. To quantify the effect of distribution shift, we measure the tree distance from each of the 1k novel concepts as the distance to its nearest neighbor among the 1k training concepts in ImageNet1k. In Supp. Fig. 1**c** we plot the average few-shot learning accuracy as a function of tree distance to the training set. Few-shot learning accuracy degrades slightly with distance from the training set, but the effect is not dramatic.

## Notes

### Competing Interest Statement

The authors have declared no competing interest.

